# RNPP-type quorum sensing regulates solvent formation and sporulation in *Clostridium acetobutylicum*

**DOI:** 10.1101/106666

**Authors:** Ann-Kathrin Kotte, Oliver Severn, Zak Bean, Katrin Schwarz, Nigel P. Minton, Klaus Winzer

## Abstract

The strictly anaerobic bacterium *Clostridium acetobutylicum* is well known for its ability to convert sugars into organic acids and solvents, most notably the potential biofuel butanol. However, the regulation of its fermentation metabolism, in particular the shift from acid to solvent production, remains poorly understood. The aim of this study was to investigate whether cell-cell communication plays a role in controlling the timing of this shift or the extent of solvent formation. Analysis of the available *C. acetobutylicum* genome sequences revealed the presence of eight putative RNPP-type quorum sensing systems, here designated *qssA* to *qssH*, each consisting of RNPP-type regulator gene followed by a small open reading frame encoding a putative signalling peptide precursor. The identified regulator and signal peptide precursor genes were designated *qsrA* to *qsrH* and *qspA* to *qspH*, respectively. Triplicate regulator mutants were generated in strain ATCC 824 for each of the eight systems and screened for phenotypic changes. The *qsrB* mutants showed increased solvent formation during early solventogenesis and hence the QssB system was selected for further characterisation. Overexpression of *qsrB* severely reduced solvent and endospore formation and this effect could be overcome by adding short synthetic peptides to the culture medium representing a specific region of the QspB signalling peptide precursor. In addition, overexpression of *qspB* increased the production of acetone and butanol and the initial (48-hour) titre of heat-resistant endospores. Together, these findings establish a role for QssB quorum sensing in the regulation of early solventogenesis and sporulation in *C. acetobutylicum*.

## INTRODUCTION

The strictly anaerobic bacterium *Clostridium acetobutylicum* is well known for its ability to convert sugars and starches into organic acids and solvents (1, 2). During the first half of the last century, the organism was used for the large scale industrial production of acetone and butanol, but the classical ABE (acetone-butanol-ethanol) fermentation process is not currently economically viable (3). Thus, considerable efforts have been devoted to improving the organism’s performance through metabolic engineering (4). However, our understanding of the organism’s physiology and metabolism, in particular the mechanisms that govern timing and extent of solvent formation, is still limited (5, 6).

In a typical *C. acetobutylicum* batch culture, acid and solvent metabolism are associated with different growth phases. During the exponential phase, a typical butyrate fermentation is carried out which allows the bacterium to maximise ATP generation. At this stage, butyric and acetic acid, as well as hydrogen and carbon dioxide, are the main fermentation products, with the former two accumulating in the culture medium. However, the increasing concentration of these short chain fatty acids poses a problem for the cells as the pH of the medium decreases and un-dissociated acids diffuse back into the cells. To avoid collapse of the proton motive force, *C. acetobutylicum* shifts its metabolism to solvent formation. In batch culture, this shift usually occurs during the transition to stationary phase and is accompanied by the partial uptake of the previously produced acids, resulting in a pH increase. These acids, together with the remaining sugars, are then converted to butanol, acetone, and ethanol (1, 2). However, solvents at high concentrations, in particular butanol, are toxic to the cells, too. The metabolic switch to solvent formation therefore leads to the initiation of yet another survival strategy: the formation of heat-resistant endospores. After solvent formation has been initiated and after cells have entered stationary phase, an intracellular starch-like storage compound termed granulose is transitorily formed and accumulates in the cytoplasm (2).

The regulatory mechanisms that govern acid and solvent metabolism, as well as sporulation, are subject to intensive research. The global transcriptional regulator Spo0A is known to be required for high solvent production in solventogenic *Clostridium* sp. and is also essential for the initiation of sporulation (7, 8). Although solvent formation in *spoA* mutants is still induced during the transition to stationary phase, the levels of acetone and butanol produced are drastically reduced (8, 9). Other regulators implied in the regulation of solventogenesis are the global regulator CodY, a small regulatory RNA, *solB*, and the catabolite control protein CcpA (10–13). The latter was shown to positively regulate the *sol* operon, which encodes key genes required for acetone and butanol formation (12).

Presumably, transcription of several solvent genes is strongly activated by the binding of phosphorylated Spo0A to specific sites, 0A boxes, which are present in the promoter regions of these genes (8, 9) Responsible for the Spo0A phosphorylation state are three orphan histidine kinases as well as an intracellular kinase-like protein which, however, acts as a phosphatase (14). Unfortunately, none of the signals or cues activating or inhibiting these proteins is currently known, although intracellular accumulation of butyrylphosphate has been proposed as a possible physiological signal and Spo0A phosphodonor (15). So while the general conditions for solventogenesis are well established, and considerable progress has been made in unravelling at least some of the regulatory networks involved, we are still largely ignorant of the cues and signals which ultimately control initiation and extent of solvent formation, and of the pathways through which they mediate their effects.

We recently proposed that quorum sensing systems may be operational in *C. acetobutylicum* and may play a role in regulating solvent metabolism (16). Quorum sensing is a mechanism of cell-cell communication that relies on small, diffusible signal molecules often referred to as autoinducers. These molecules are secreted during growth, accumulate in the extracellular environment, and allow bacteria to coordinate gene expression with cell population density. In the *Firmicutes*, quorum sensing systems are usually based on secreted autoinducing peptides (AIPs) which can be linear or cyclic, and sometimes contain post-translational modifications (17, 18). Relatively little is known about the operation of such systems in clostridial species, but we have previously hypothesised (16) that quorum sensing might play a role in the regulation of solventogenesis based on the following considerations. First, for the solventogenic *Clostridium sacharoperbutylacetonicum* an as yet unidentified, self-generated signal present in the supernatant of wild type cultures was capable of inducing solvent formation in a ‘low-solvent’ mutant (19). Second, in *C. acetobutylicum* and related “high solvent” producers such as *Clostridium beijernickii*, solventogenesis during batch culture growth is usually initiated at high cell densities. Third, genome sequencing has revealed a large number of putative quorum sensing systems within the Genus *Clostridium*, including all currently sequenced solvent-producing species (20; and unpublished data from this laboratory). Furthermore, a novel polyketide signal, clostrienose, has recently been shown to affect granulose formation, sporulation and, to a smaller degree, butanol production (21).

We therefore investigated the role of a putative *agr*-type quorum sensing system in *C. acetobutylicum* ATCC 824, which we showed to be functional and involved in the regulation of sporulation and the production of granulose (16). However, the formation of organic acids and solvents from glucose was unaffected in mutants in which this system had been inactivated, suggesting that it played no role in the regulation of fermentation metabolism. Interestingly, however, the *C. acetobutylicum* genome has also been reported to contain two potential quorum sensing systems that bear similarity to the *rap*-*phr* systems present in *Bacillus subtilis* (22). In *Bacillus subtilis*, the *rap* genes encode a conserved group of regulatory phosphatases acting on phosphorylated response regulators, whereas the *phr* genes encode the precursors of short, linear signalling peptides that can bind to and inhibit the Rap proteins. The Rap proteins are part of the RNPP-type family of quorum sensing regulators, which derived its name from its best studied members, i.e. <u>R</u>ap, <u>N</u>prR, <u>P</u>lcR, and <u>P</u>rgX, and is characterised by the presence of tetratricopeptide repeat (TPR) domains responsible for promoting protein-protein interaction. The family comprises all currently known Gram-positive cytoplasmic quorum sensing regulators which directly bind to their cognate signalling peptide (23, 24). The signalling peptide is derived from the C-terminal part of the original pre-propeptide which is exported and further processed to its mature form. The mature signalling peptide is transported back into the cells by oligopeptide permeases belonging to the family of ATP-binding cassette (ABC) transporters. Thus, the uptake of these signalling molecules is an ATP-consuming process (22). Apart from the Rap proteins, all other currently identified RNPP-type regulators, including the two proposed *C. acetobutylicum* homologues (CA_C186 and CA_C3694), possess helix-turn-helix (HTH) motifs and are either known or likely to be transcriptional regulators that become activated or inhibited upon binding their cognate signal peptide (24).

Here, we report the discovery and mutational screening of eight putative RNPP-type quorum sensing systems in *C. acetobutylicum* ATCC 824 and the more detailed characterisation of one of these systems, QssB.

## RESULTS

### Identification of eight putative RNPP-type quorum sensing systems in *C. acetobutylicum*

Using the two previously identified *C. acetobutylicum* homologues (22) and other experimentally confirmed HTH-containing RNPP-type regulators such as PlcR and NprR in blastp searches, a total of eleven putative RNPP-type regulators genes were identified in published *C. acetobutylicum* genomes (strains ATCC 824, DSM 1731, and EA 2018). The locus tags for these genes in the ATCC 824 strain were CA_C0186, CA_C0324, CA_C0957, CA_C0958, CA_C1043, CA_C1214, CA_C1949, CA_C2490, CA_C3694, CA_P0040 and CA_P0149. Most of them were annotated as either hypothetical proteins or regulators of the Xre family containing TPR domains, with CA_C0186 and CA_C3694 representing the previously identified homologues (22). In the current version of the ATCC 824 genome (NC-_003030.1), CA_C3694 is annotated as a pseudogene in which the HTH motif and TPR domain-encoding parts of an RNPP-type regulator are separated by a stop codon. However, in the genomes of the DSM1731 and EA 2018 strains, these domains are encoded by a single gene. We therefore compared the published ATCC 824 sequence to that obtained for our version of this strain (25) and also found the two domains to be encoded by a single gene. In the original ATCC 824 sequence, the insertion of a guanine at position 357 had shifted the reading frame so that a stop codon appeared to terminate CA_C3694 translation after 360 bp. The corrected sequence was identical to that in the EA2018 and DSM 1731 strains (26, 27).

To establish a putative role in quorum sensing, the regions flanking the above regulator genes were analysed for the presence of short ORFs encoding putative quorum sensing peptide precursors. For established RNPP-type systems, these precursors consist of a positively charged N-terminus, followed by a hydrophobic region (together forming a signal peptide sequence, required for peptide export) and a C-terminal part containing the actual autoinducing peptide (22). Short ORFs fulfilling the above criteria could be identified downstream of the CA_C0186, CA_C0324, CA_C1043, CA_C1214, CA_C2490, CA_C3695/CA_C3694, CA_P0040 and CA_P0149 (Fig. 1A). Only one of these ORFs (CA_C3693) was annotated in the ATCC 824 genome sequence. The eight aforementioned regulator genes were therefore designated quorum sensing regulators A to H (*qsrA* to *qsrH*), and their cognate quorum sensing peptide (Qsp)-encoding genes *qspA* to *qspH*, respectively. Together they were referred to as quorum sensing systems A to H (QssA to QssH).

**Figure 1.**
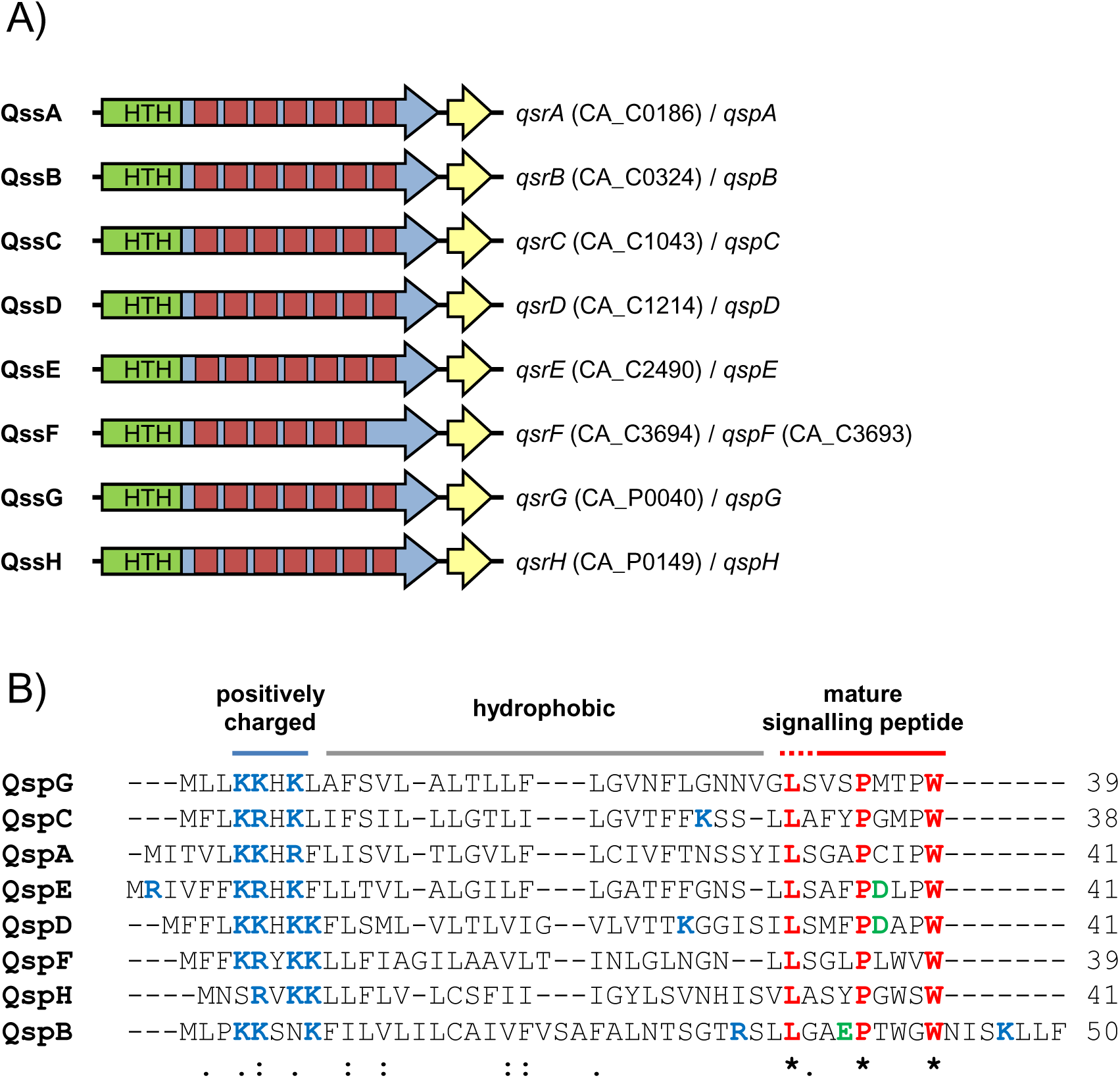
Schematic representation of *C. acetobutylicum* RNPP quorum sensing gene clusters (A) and alignment of putative signalling peptide precursor sequences (B). (A) Eight RNPP quorum sensing gene clusters have been identified (QssA to QssH), each encoding an RNPP-type regulator (QsrA to QspH, large arrows) and a signalling peptide precursor (QspA to QspH, short yellow arrows). The locus tags for each system are provided, where available. Regions encoding a helix-turn-helix motif (HTH, green) and tetratricopeptide repeat domains (red) are indicated. (B) The Clustal Omega amino acid sequence alignment shows the eight predicted Qsp proteins. Red font indicates amino acids that are 100% conserved; blue and green fonts indicate positively (K, R) and negatively charged (D, E) amino acids, respectively. Identical (*), conserved (:), and semi-conserved substitutions (.) are shown. Numbers indicate the length of the different precursor proteins. Positively charged, hydrophobic, and predicted signalling peptide-encoding regions are indicated with blue, grey, and red lines, respectively.

Comparison of the identified Qsp sequences revealed that they were of similar length but low overall sequence similarity (Fig. 1B). However, three amino acids at the C-terminal end were conserved in all Qsp proteins: a leucine, a proline and a tryptophan. The latter formed the C-terminal amino acid in all putative Qsp proteins, with the exception of QspB, which was extended by an additional seven amino acids.

The deduced Qsr sequences were also of similar length and showed low overall similarity, apart from N-terminal region, which contained a Xre-type HTH motif. Using TPRpred (28), the remaining parts of the Qsr proteins were predicted to each possess 7 TPR domains, with the exception of QsrF, for which only six putative domains were detected. Comparison with other HTH-containing RNPP-type regulators revealed low identity and similarity values, with very few conserved amino acid positions, again mainly positioned in the predicted HTH-domains of the proteins (data not shown).

Analysis of available genome sequences revealed the presence of similar systems in other members of the class *Clostridia,* notably the solventogenic *C. saccharoperbutylacetonicum* strain N1-4 the genome of which was found to contain five putative RNPP-type gene clusters (see Fig. S1 in the supplemental material). However, no such systems (i.e. containing both regulator and peptide) could be identified in the closely related *C. beijerinckii*.

### Insertional inactivation of *qsr* genes using ClosTron technology

Using ClosTron technology (2), all eight identified *qsr* genes were insertionally inactivated in the ATCC 824 strain. Correct insertion of *ermB*-carrying introns into the target genes was confirmed by PCR screens and sequencing of the obtained PCR products as described previously (16).

The chosen ClosTron insertion sites were located within the putative HTH-encoding region of the *qsr* genes (see Methods), thus ensuring that no active DNA-binding proteins could be formed. For each gene, at least three independently derived ClosTron clones were isolated and further characterised. This was done to avoid accidental isolation of mutant clones carrying undesired second site mutations: solventogenic *Clostridium* species including *C. acetobutylicum* are known to spontaneously ‘degenerate’, resulting in strains with a reduced or abolished capacity to form solvents and heat-resistant endospores (30, 31). A preliminary analysis revealed that one of the independently obtained *qsrC* mutant clones differed phenotypically from the other three and showed signs of degeneration (data not shown). This clone was therefore excluded from further phenotypic screening.

### Phenotypic screening of *qsr* mutants to identify systems of interest

The obtained *qsr* mutants were phenotypically characterised with respect to growth, colony morphology, starch degradation, granulose formation, sporulation, and solvent formation.

When cultured in CBMS broth, several mutants showed minor differences in their growth kinetics when compared to the wild type (see Fig. S2 in supplemental material). Under the conditions employed, wild type cultures reached an OD_600_ of 2.6 after 9 h, followed by a transient OD_600_ decrease to 1.4 (24 h) before reaching the final maximum OD_600_ of 3.1 (48 h). Concurrent with the transient decrease in OD_600_, the cultures began to appear more viscous. Similar profiles were observed for the mutant strains, although *qsrF* and *qsrG* mutants reached lower final ODs after 48 h (1.92 and 1.03, respectively), *qsrC* and *qsrD* mutants grew more slowly, and *qsrB* mutant cultures showed the transitory OD_600_ decrease and viscosity increase already after 12 h. The low final OD_600_ of the *qsrG* mutant presumably reflected the strain’s tendency to form large cell aggregates.

The ability to degrade starch was not affected in any of the mutants and granulose formation appeared similar to the wild type (data not shown). Interestingly, however, after 24 h of growth on CGM plates the *qsrB* mutants were observed as forming considerably larger colonies (2.0 mm) when compared to the wild type (1.3 mm), a difference that was statistically significant (p<0.00001; Table 1).

**Table 1.** Colony size of *Clostridium acetobutylicum* parents strain and *qsrB* mutants on CGM medium after 24 h.

| Strain | Mean colony size<br>in mm ±SD (n=20) |
| --- | --- |
| *C. acetobutylicum* ATCC 824 | 1.3 ±0.49 |
| *C. acetobutylicum qsrB::CTermB* | 2.0 ±0.24***** |
| *C. acetobutylicum* ATCC 824 pMTL 85141 | 1.2 ±0.32 |
| *C. acetobutylicum qsrB::CTermB* pMTL85141 | 1.5 ±0.35** |
| *C. acetobutylicum qsrB::CTermB* pMTL85141-*qsrB* | 1.0 ±0.28<sup>NS</sup> |

<sup>NS</sup> not significantly different to the vector carrying wild type; \*\*significantly different to the vector carrying wild type at p<0.01; \*\*\*\*\*significantly different to the vector-free wild type control at p<0.00001; n=20.

Microscopic examination of CBMS grown cultures revealed no noticeable changes in the number of endospores formed by *qsr* mutants when compared to the wild type. However, a significant 3-fold reduction (p=0.036) was observed for *qsrG* mutants when a more quantitative procedure was used, i.e. when the number of heat-resistant spores in a given culture volume was determined after 7 days of culture (Table S1; more precisely, this procedure quantifies the number of heat-resistant colony forming units (cfu) as a measure for spores that can germinate and grow after heat treatment at 80 °C for 10 min).

The ability of *qsr* mutants to form butanol, acetone, and ethanol was also assessed during early (24 h) and late (120 h) solventogenesis. According to their butanol production profiles (Fig. S3), *qsr* mutants could be grouped into four categories: (i) early and late butanol titres similar to the wild type: *qsrC* and *qsrD* mutants; (ii) increased butanol titres during early solventogenesis: *qsrB* mutants; (iii) decreased butanol titres during early solventogenesis: *qsrA* and *qsrE* mutants; (iv) decreased butanol titres during late solventogenesis: *qsrF*, *qsrG*, *qsrH*. Generally, changes in butanol titers were mirrored by the corresponding acetone concentrations. However, this was not always the case for ethanol. For instance, final (120 h) ethanol titres were significantly increased for the *qsrB* and *qsrE* mutants and early (24 h) titres in *qsrA* and *qsrE* mutants were comparable to those of the wild type (Fig. S3).

### QsrB represses solvent formation

Following the initial phenotypic screening, the *qsrB* mutants were selected for a more thorough characterisation as they exhibited a number of relevant phenotypic changes including growth profile, colony size/morphology, and solvent production. Particularly relevant from a biotechnological perspective was the increased production of butanol during early solventogenesis. More detailed fermentation profiles were therefore generated, with samples taken at regular intervals during a 120 h growth experiment. These profiles confirmed the increased production of solvents during early solventogenesis and also revealed that, after entry into stationary phase, *qsrB* cultures contained lower concentrations of butyric and acetic acid (Fig. S4 in supplemental material). To obtain ultimate proof that *qsrB* inactivation was responsible for the observed phenotypes, the obtained *qsrB* mutants were genetically complemented. *qsrB* under control of its native promoter was cloned into the modular shuttle vector pMTL85141 (32) and the resulting pMTL85141-*qsrB* vector was used to transform the *qsrB* mutant strains via electroporation. As a control, unmodified pMTL85141 was also introduced into both wild type and *qsrB* mutant strains. Indeed, complementation with plasmid-based *qsrB,* but not the empty shuttle vector, reversed the effects of *qsrB* inactivation, i.e. it reduced the production of all three solvents, increased the production of acetic and butyric acid, and reduced colony size (Fig. 2 and Table 1). However, while colony size was restored to approximately wild type levels, the metabolic changes resulting from the complementation were more drastic, i.e. solvent production by the complemented *qsrB* mutants was significantly lower, and acid production markedly higher, than observed for the wild type. It was hypothesised that this was caused by the presence of multiple *qsrB* copies in the complemented mutants. Indeed, when these experiments were repeated using the overexpression vector pMTL85143, which carries a strong constitutive ferredoxin gene promoter to drive the expression of the inserted *qsrB* gene, very similar results were obtained (data not shown). Expression of *qsrB* in the wild type using the pMTL85141-*qsrB* and pMTL85143-*qsrB* plasmids yielded fermentation profiles similar to the ones observed for the complemented *qsrB* mutant, with increased production of acids and considerably reduced solvent formation (Table 2).

**Figure 2.**
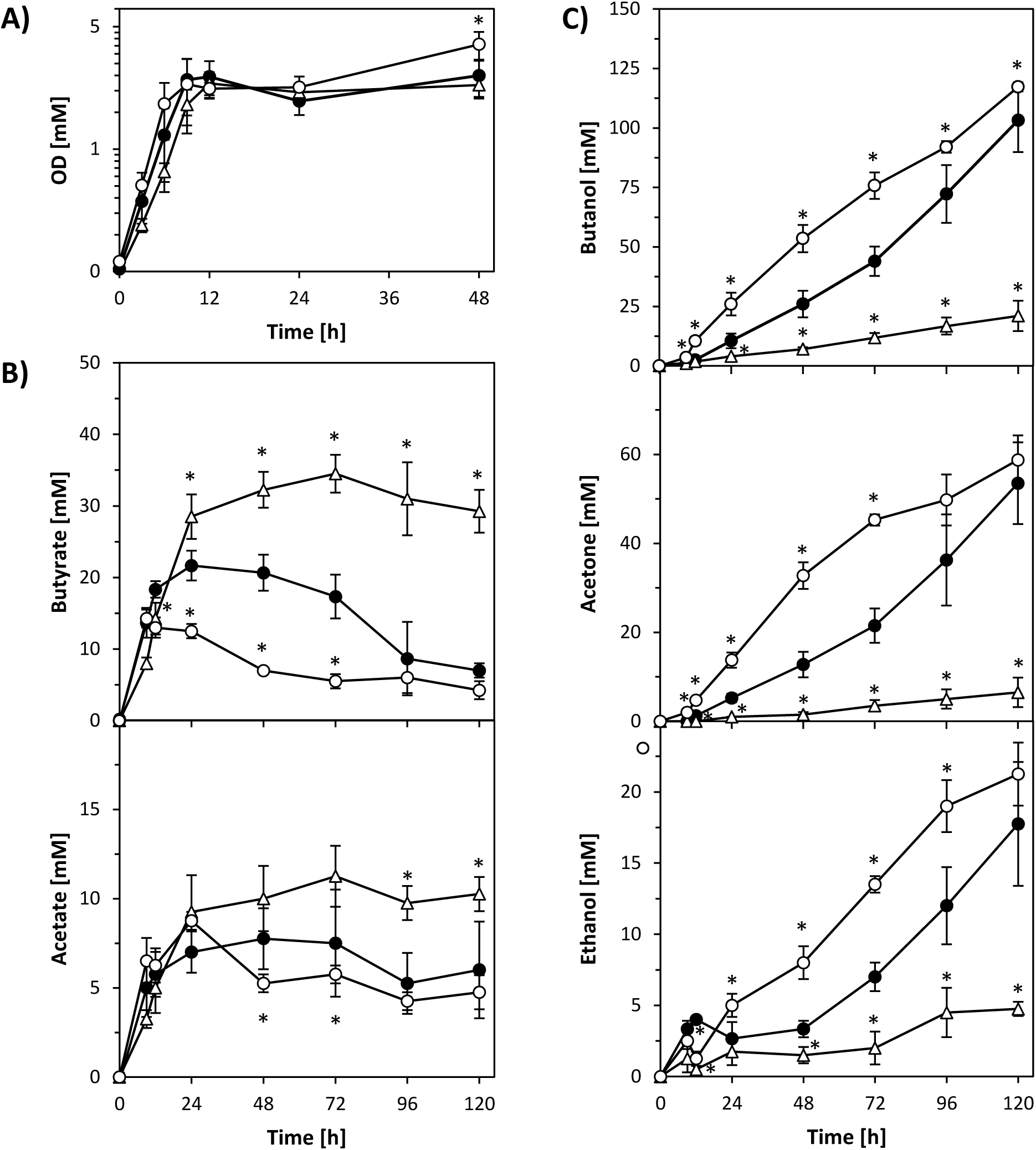
Fermentation profile of *C. acetobutylicum* wild type, *qsrB* mutants and genetically complemented *qsrB* mutants. Growth (A) and production of acids (B) and solvents (C) were compared for the ATCC 824 parent strain containing the empty pMTL85141 vector (closed circles), the *qsrB* mutants containing the empty pMTL85141 vector (open circles), and the *qsrB* mutants containing the complementation plasmid pMTL85141-*qsrB* (open triangles). Data represent the mean of four independent CBMS cultures with error bars indicating the standard deviation. Significant differences (p≤0.05) compared to the wild type are indicated by an asterisk next to the relevant data point.

**TABLE 2.** Effect of *qsrB* overexpression on the fermentation product profile of wild type *C. acetobutylicum* ATCC 824 after 120 h growth in CBMS.

| Product<br>[mM] | *C. acetobutylicum* |  |  |  |
| --- | --- | --- | --- | --- |
|  | pMTL85141 | pMTL85141-*qsrB* | pMTL85143 | pMTL85143-*qsrB* |
| Butyrate | 7 ±1 | 36 ±3***** | 13 ±13 | 48 ±15***** |
| Acetate | 6 ±3 | 14 ±1** | 25 ±18 | 35 ±16* |
| Butanol | 103 ±15 | 15 ±3***** | 86 ±23 | 9 ±4*** |
| Acetone | 54 ±4 | 4 ±1***** | 38 ±18 | 1 ±1** |
| Ethanol | 18 ±4 | 4 ±3** | 15 ±9 | 1 ±1* |

\*, \*\*, \*\*\* and \*\*\*\*\* indicate significant differences to the vector carrying wild type at p<0.05, p<0.01, p<0.001, and p<0.0001, respectively; n=3.

### Overexpression of *qsrB* reduces spore formation

Given the drastic effects that *qsrB* carrying plasmids had on acid and solvent formation, the number of heat-resistant endospores formed by the complemented *qsrB* mutants and *qsrB* overexpressing wild type were also assessed. These experiments revealed that in the presence of pMTL85141-*qsrB* both strains showed strongly reduced spore production (Fig. 3). Furthermore, while after 120 h and 168 h there was no statistically significant difference in spore counts between wild type and *qsrB* mutants that both carried the empty pMTL85141 control plasmid, the latter reached final spore levels earlier than the wild type. This suggested earlier or more rapid sporulation in the absence of *qsrB*.

**Figure 3.**
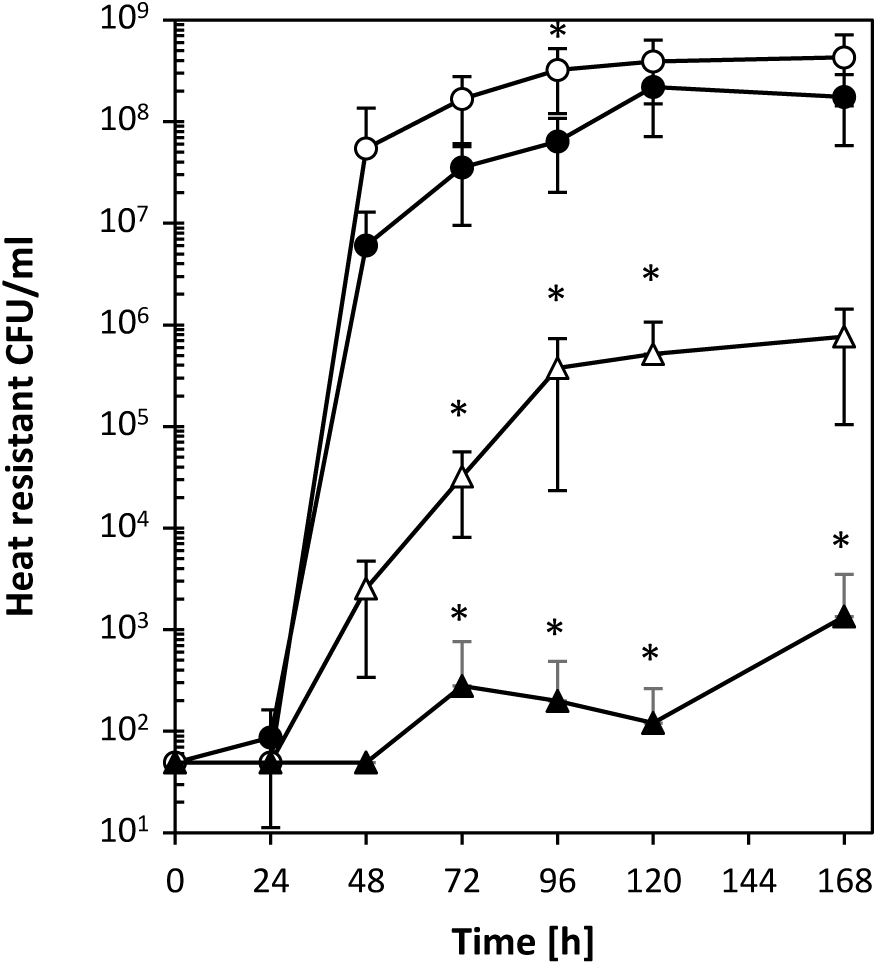
Effect of *qsrB* deletion and overexpression on sporulation. The ability to sporulate in CBMS was assessed for the ATCC 824 parent strain containing the empty pMTL85141 vector (closed circles), the *qsrB* mutants containing the empty pMTL85141 vector (open circles), the *qsrB* mutants containing the pMTL85141-*qsrB* complementation plasmid (open triangles), and the ATCC 824 parent strain containing the pMTL85141-*qsrB* complementation plasmid (closed triangles). Sporulation efficiencies were assessed by determining the number of heat-resistant endospores produced at the indicated time points. Data represent the mean of four independent CBMS cultures with error bars indicating the standard deviation. Only the upper half of the error bar is shown in cases where the lower half extends beyond 10^-1^. Significant differences (p≤0.05) compared to the vector-carrying wild type and vector-carrying *qsrB* mutant are indicated by an asterisk next to the relevant data point.

### Generation and characterisation of *qspB* mutants

Based on the above results it appeared likely that *qsrB*-based quorum sensing contributes to the regulation of solvent formation and sporulation in *C. acetobutylicum*. To test this hypothesis, the role of the putative signalling peptide-encoding *qspB* gene, located downstream of *qsrB*, was investigated. Three independent *qspB* ClosTron mutants were generated and confirmed as described above for the *qsrB* mutants. While colony size, granulose formation, and final spore levels, were comparable to the wild type, *qspB* mutant cultures showed reduced levels of acetone and butanol during late stationary phase, i.e. after 72 h to 96 h. However, final (120 h) levels were comparable to the wild type (data not shown). Introduction of the aforementioned shuttle vectors (pMTL85141 and pMTL85143; without insert) into *qspB* mutants and wild type abolished the observed differences and led to indistinguishable solvent profiles (Fig. 4). Thus, conclusive genetic complementation experiments could not be conducted. However, when the *qspB* overexpression plasmid pMTL85143-*qspB* was introduced into both wild type and *qspB-* deficient strains, solvent production increased significantly and butyrate concentrations during stationary phase were considerably lower than in the control strains carrying the empty pMTL85143 plasmid. Acetate production, however, remained largely unchanged (Fig 4B, 4C). Overexpression of *qspB* also increased colony size and lead to an earlier increase in heat-resistant colonies, although final spore levels appeared to be similar to the wild type vector control (Fig. 5).

**Figure 4.**
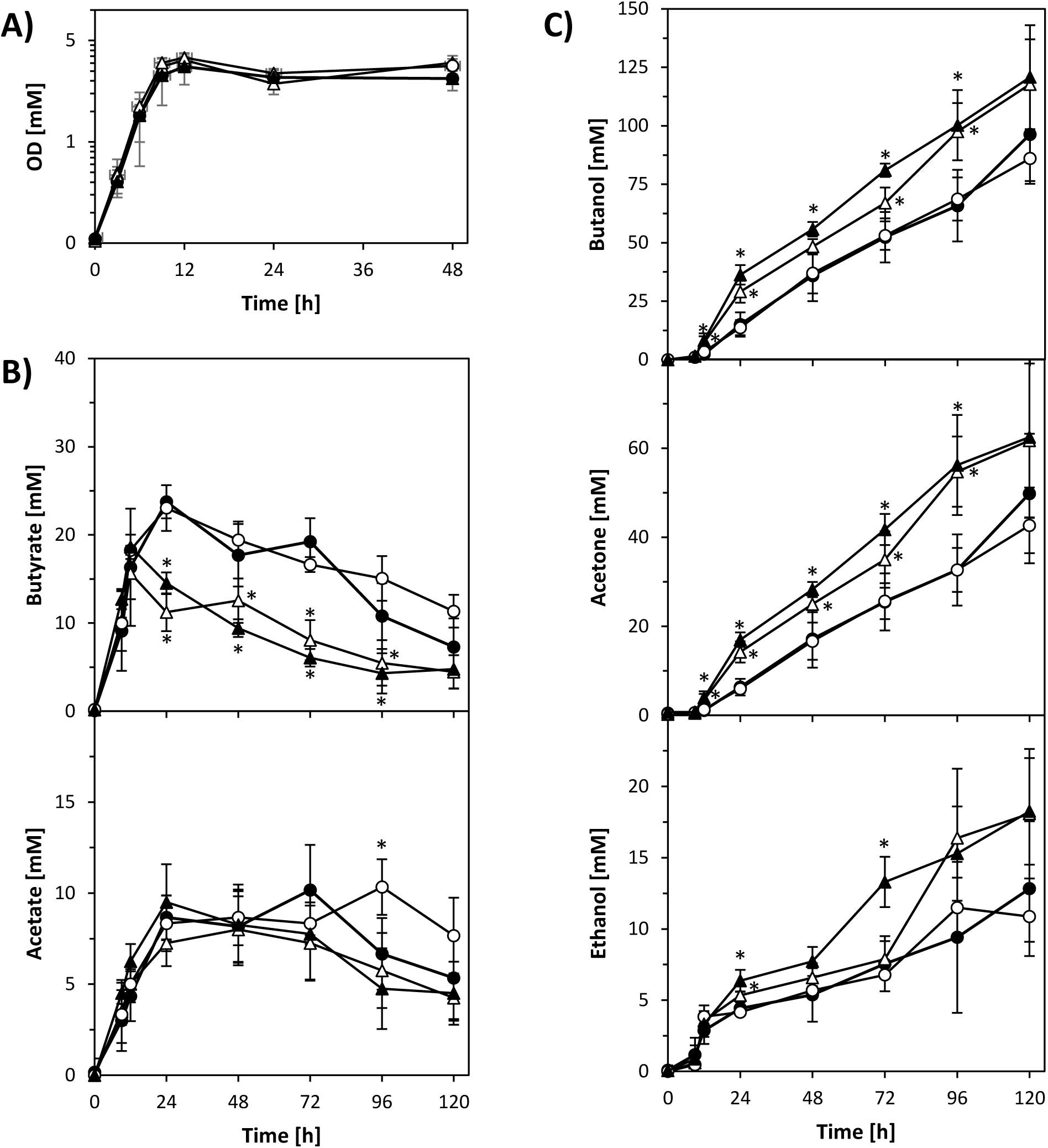
Fermentation profiles of *qspB*-overexpressing *C. acetobutylicum* wild type and *qspB* mutants. Growth (A) and production of acids (B) and solvents (C) were compared for the ATCC 824 parent strain (closed symbols) and *qspB* mutant (open symbols) containing either the empty pMTL85143 vector (circles) or the overexpression plasmid pMTL85143-*qspB* (triangles). Data represent the mean of four independent CBMS cultures with error bars indicating the standard deviation. Significant differences (p≤ 0.05) compared to the vector controls are indicated by an asterisk next to the relevant data point.

**Figure 5.**
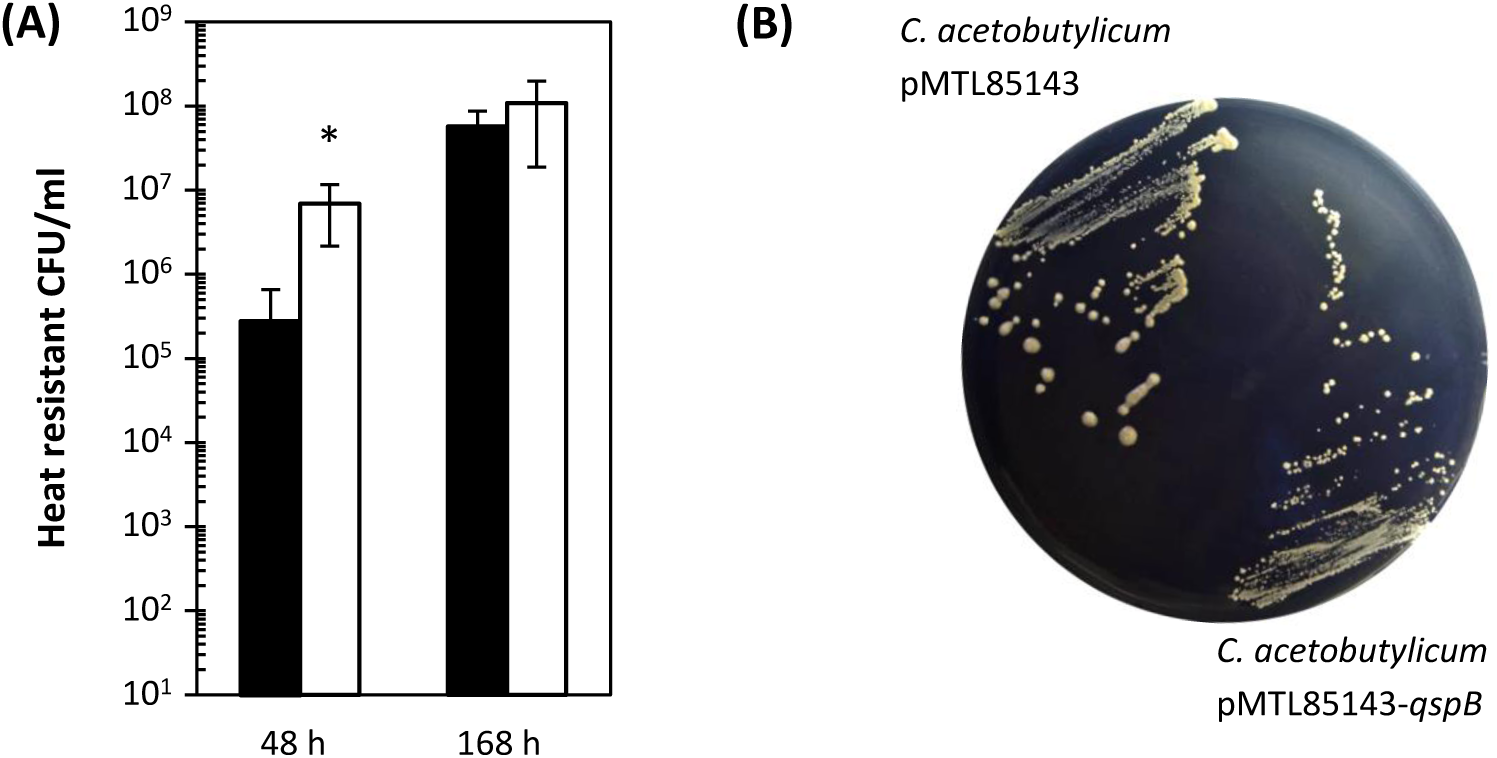
Effect of *qspB* overexpression on sporulation and colony size. (A) The ability to sporulate in CBMS was assessed for the *C. acetobutylicum* ATCC 824 parent strain containing the empty pMTL85143 vector (black bars) and the overexpression plasmid pMTL85143-*qspB* (white bars). Data represent the mean of four independent cultures with error bars indicating the standard deviation. Significant differences (p≤ 0.05) compared to the vector controls are indicated by an asterisk next to the relevant measurement. (B) 5-day old colonies of *C. acetobutylicum* carrying the empty pMTL85143 vector (left) and overexpression plasmid pMTL85143-*qspB*, respectively, on CGM agar.

### *qspB*-encoded peptide fragments counteract QsrB-mediated repression of solventogenesis and sporulation

The similar phenotypes observed for *qsrB*-knockout and *qspB*-overexpressing strains suggested that either QspB or a QspB-derived quorum sensing peptide might act to inhibit QsrB activity. To test the latter hypothesis, thirteen peptide variants were synthesised, varying in length between 6 and 20 amino acids and covering various parts of the C-terminal region of QspB. These were then tested for their ability to restore butanol production in the *qsrB* overexpressing strain *C. acetobutylicum* pMTL85141-*qsrB*. Cultures of this strain were supplemented with individual synthetic peptides at a final concentration of 10 μM and assayed for butanol formation after 120 h. Several of the exogenously added peptides were capable of restoring high-level butanol formation, whereas others had no discernible effect (Fig. S5 in the supplemental material). The latter group comprised all peptides terminating at amino acid 38 of the QspB sequence or starting at position 39, suggesting that the region conferring activity included amino acids upstream and downstream of these positions.

Based on these finding, additional peptide variants were designed, synthesised to a purity of >95% and similarly tested. Interestingly, exogenous addition of QspB7, a peptide comprising only seven amino acids (AEPTWGW) and matching positions 37-43 of the QspB precursor, was capable of fully restoring butanol production in the *qsrB* overexpressing *C. acetobutylicum* pMTL85143-*qsrB* strain (Fig. 6). The QspB7 sequence contained two of the three conserved amino acids present at the C-terminal end of all *C. acetobutylicum* Qsp proteins, i.e. proline and tryptophan (Fig. 1B). QspB-derived peptides capable of restoring high-level butanol formation were also found to dramatically increase acetone and decrease acid production when added to the *qsrB* overexpressing strain (Fig. 6A, 6B). Furthermore, these peptides also restored high levels of sporulation (Fig. 6C)

**Figure 6.**
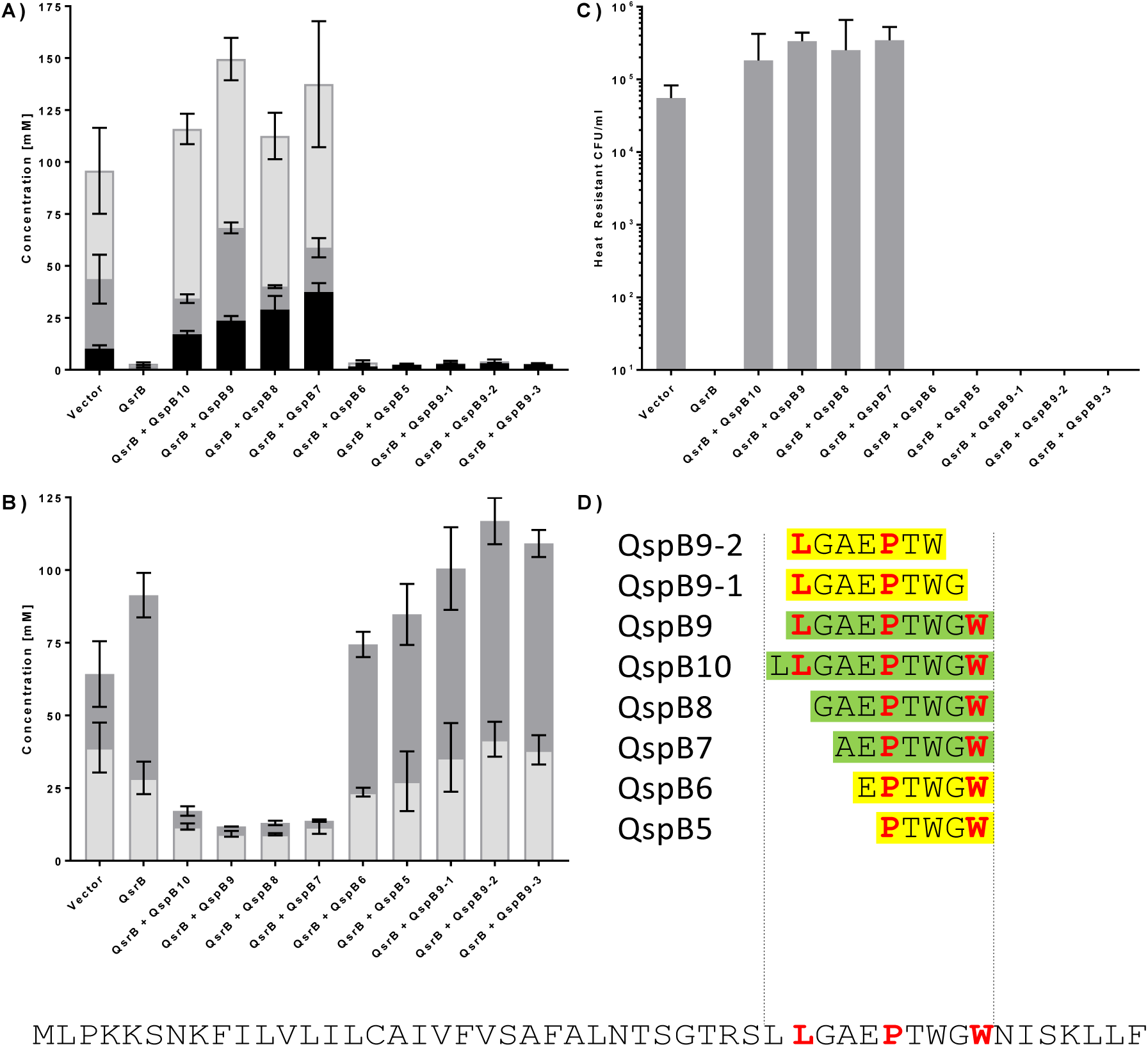
Synthetic peptides alleviate *qsrB*-mediated repression of solvent formation and sporulation. (A) Solvent titres: butanol (light grey), acetone (dark grey), ethanol (black). (B) Acid titres: butyrate (dark grey), acetate (light grey). (C) Spore titres (heat-resistant CFU). Synthetic peptides (D) were dissolved in DMSO and individually added to CBMS cultures of *C. acetobutylicum* pMTL85143-*qsrB* after 4 h of growth to a final concentration of 10 μM. DMSO-only controls were performed for *C. acetobutylicum* pMTL85143 and *C. acetobutylicum* pMTL85143-*qsrB,* respectively. Cultures were grown for 5 days prior to analysis. *C. acetobutylicum* pMTL85143 vector control (Vector); *C. acetobutylicum* pMTL85143-*qsrB* cultures (QsrB). Presence of specific synthetic peptides as shown in (D) is indicated (+ QspB). The complete QspB sequence is given at the bottom with the three conserved amino acid positions in the C-terminal region (leucine, proline, tryptophan) indicated by bold red lettering. Data represent the mean of three independent cultures with error bars indicating the standard deviation.

## DISCUSSION

For many years, the precise molecular signals and mechanisms that trigger and control solvent formation in clostridia have remained elusive. Here we show that RNPP-type quorum sensing is one of the contributing factors in *C. acetobutylicum* and suggest that this may also apply to other solventogenic species.

RNPP-type systems consist of two components, a TPR domain-containing regulator protein and a secreted linear signalling peptide, which following uptake will bind to and either activate or inactivate the regulator (24). Examples include the *B. subtilis* Rap-Phr systems involved in the regulation of competence and sporulation and the <u>P</u>lcR-PapR systems of the *Bacillus cereus* group, which control toxins and other genes important for environmental adaptation (22, 33). This study set out to examine the roles of putative RNPP-type quorum sensing systems in *C. acetobutylicum* and, more specifically, whether this type of communication plays a role in the regulation of acid and solvent metabolism.

Bioinformatic analysis revealed the presence of at least eight such systems in several sequenced strains of this bacterium, including two that had previously been proposed (22). To identify systems of interest, all eight were subjected to ClosTron mutagenesis resulting in disruption of the postulated DNA-binding HTH motifs encoded in the 5’ regions of the respective *qsr* genes. Like all mutagenesis procedures that involve the frequent re-streaking of colonies, the ClosTron method carries an inherent risk of enriching for and isolating degenerate mutant strains (16). At least three independent mutant clones were therefore generated for each *qsr* gene and checked for phenotypic consistency, thus ensuring that observed phenotypic changes were caused by ClosTron insertion rather than random second site mutations.

As a detailed analysis of all eight systems was beyond the scope of this study, a general phenotypic screen was carried out to identify mutants of interest, i.e. those showing clear phenotypic differences in particular with relation to solvent metabolism. Intriguingly, inactivation of seven of these systems resulted in changes to the observed solvent profiles, although further studies will be necessary to establish whether these systems are directly involved in the regulation of solvent genes, or whether these are indirect effects resulting from other changes in growth, physiology and metabolic activity. Based on this initial screen, QssB was selected for or a more detailed characterisation, as it was the only system whose inactivation increased solvent formation and also affected multiple other phenotypes.

Taken together, our mutational analyses suggest that QssB plays a regulatory role during early solvent formation, controlling its extent and, potentially, precise timing. The data are consistent with QsrB acting as a repressor that is inactivated upon binding to its cognate QspB-derived signalling peptide. Alternatively, QsrB might exert its effects by positively regulating an unknown repressor of solvent formation. Both scenarios are supported by the finding that (i) solvent formation was increased by *qsrB* inactivation and strongly decreased by *qsrB* overexpression; (ii) overexpression of *qsrB* and *qspB* had opposing effects on solvent formation, sporulation and colony size; and (iii) repression of solvent formation and sporulation in *qsrB* overexpressing cells could be overcome by adding short synthetic QspB-derived peptides to the culture medium. By contrast, the strong reduction of endospore formation seen for the *qsrB* overexpressing strains might be an indirect consequence of the drastically reduced solvent metabolism.

It is unclear why inactivation of the signalling peptide encoding *qspB* gene had only limited effects during early solventogenesis, as under these conditions QsrB might be expected to repress, directly or indirectly, solvent formation in the absence of its cognate signalling peptide. An intriguing possibility could be that QsrB responds to more than one signalling peptide, i.e. lack of QspB may be compensated for by signalling peptides produced by the other quorum sensing systems. This may also explain why the sporulation profile of the *qspB* mutant was so similar to that of the wild type. The fact that *qspB* overexpression resulted in a small increase in sporulation may indicate that, under the employed culture conditions, wild type signal molecule concentrations were not sufficiently high to completely inactivate the QsrB regulator at the time when this process was induced, consistent with the slight increase in sporulation observed for *qsrB* mutants.

An interesting question is why quorum sensing control of solvent formation has evolved in *C. acetobutylicum*. A possible explanation could be that coordinated, population-wide responses are required to efficiently control the rapid production of toxic acids and, perhaps, at a later stage, solvents. For an optimal response, individual cells within the population may need to sense the density of acid producing cells and this could be achieved by the extracellular accumulation of peptide signals such as those derived from QspB. Thus, before critical concentrations are irreversibly reached that lead to a fatal ‘acid crash’ (34), a population-wide decision is made to stop production and trigger a metabolic shift resulting in acid re-uptake and solvent formation. Integrated with other relevant environmental information this may enable the organism to maximise the number of cells that can enter solventogenesis and thus, eventually, sporulation to secure long-term survival. According to this hypothesis, uptake of acids and their conversion into solvents would be a social, cooperative trait, which is co-ordinately induced through quorum sensing at the appropriate population density.

It is evident from the above that sensing their density may help populations to shift from acid to solvent metabolism at an optimal stage of growth. It is less clear, however, why the organism contains such a large number of putative signalling systems, totalling ten together with the previously described *agr* (16) and clostrienose systems (21). For many bacterial species, including pathogens, no more than one or two systems have been identified, although larger numbers are present in some ubiquitous and/or metabolically versatile bacteria such as *Bacillus subtilis* or *Pseudomonas aeruginosa*) (35, 36).

It is therefore intriguing to see that two other, physiologically very similar, solvent producing *Clostridium* species also contain a large number of putative signalling systems. Our analysis of the *C. saccharoperbutylacetonicum* genome revealed the presence of five complete RNPP-type systems (Fig. S1) in addition to four potential *agr* system (not shown). By contrast, *C. beijernickii* NCIMB 8052 does not appear to contain complete RNPP-type systems, but has six putative *agr* systems. Thus, while all three species are members of the genus *Clostridium sensu stricto*, carry out very similar ABE fermentations and, in the case of *C. beijernickii* and *C. saccharoperbutylacetonicum*, are very closely related based on 16S rRNA sequence similarity, they differ considerably in terms of the cell-cell signalling systems they employ (37).

Most likely, the explanation for employing multiple signalling systems lies in the complex life cycles of these organisms which not only involve a shift in fermentation metabolism, but also endospore formation and, under certain conditions, fruiting body formation (38). These are all phenotypes for which quorum sensing control has been demonstrated in other species. Furthermore, use of multiple signals may permit ‘combinatorial communication’, enabling bacteria to adjust gene expression to both social and physicochemical properties of their environment, particularly when accumulation of the individual signal molecules differs due to differences in half-life or diffusion constants (39). Alternatively, non-combinatorial use of multiple peptide signals may simply enable cells to trigger responses at different density thresholds. To add to this complexity, the genomes of *C. acetobutylicum* and indeed all other members of the genus *Clostridium sensu stricto,* encode several orphan RNPP-type regulators that are not flanked by small, signalling peptide encoding genes. Whether genes of this type form part of a quorum sensing systems or act independently of signalling peptides remains to be seen, but there is evidence to suggest that they, too, play major regulatory roles in their respective hosts. For instance, the CA_C0957/CA_C0958 regulators identified in this study were found to be important for solventogenesis and sporulation (40) and in *Clostridium difficile*, another orphan RNPP-type regulator was recently found to repress toxin production and motility, and upregulate sporulation (41).

The precise chemical nature of the Qsp-derived peptide signals produced by *C. acetobutylicum* remains to be established. In *B. subtilis*, some of the Phr peptides are derived from the C-termini of their respective precursor peptides whereas others stem from internal regions (22). A similar situation appears to be present in *C. acetobutylicum*. Our structure activity analysis of QspB derived peptide sequences clearly showed that biological activity is associated with a short internal sequence. However, for the other seven putative Qsp homologs, sequences corresponding to this region form the C-terminal end of the precursor peptide (Fig. 1).

Whereas the Phr signals produced by *B. subtilis* are pentapeptides, the *C. acetobutylicum* QspB peptide appears to be slightly larger given that a heptapeptide was the shortest sequence for which biological activity was observed (Fig. 6). This heptapeptide, AEPTWGW, contained two of the three conserved amino acids present in the C-terminal region of all Qsp proteins, i.e. proline and tryptophan, whereas a slightly larger nonamer, LGAEPTWGW, showed similar activity but also contained the third conserved amino acid, leucine. This is similar to findings made for PlcR and its cognate heptapeptide signal, PapR, in the *B. cereus* group. Originally believed to be a pentapeptide due to its biological activity, the native PapR signal was later found to be a heptamer (42). PapR peptides from different strains of this group show some variation in the first three N-terminal residues, whereas the C-terminal parts are relatively conserved (43). Although the predicted *C. acetobutylicum* Qsp peptides show a larger degree of variation, the aforementioned proline (position −5) and tryptophan (position −1) are always present. Interestingly, the peptides produced by *B. subtilis* and the *B. cereus* group all contain charged amino acids (22, 43), whereas this is not the case for the majority of the predicted Qsp peptides. Only QspB, QspD and QspE are predicted to carry a negative charge, whereas all other Qsp peptides are highly hydrophobic. Whether and how this relates to their biological roles remains to be investigated.

In summary, we have identified multiple signalling systems in *C. acetobutylicum*, at least one of which plays a role in the regulation of solvent and spore formation. Signal molecule accumulation appears to be an important parameter that, together with other environmental and internal stimuli, is sensed and integrated by a complex regulatory network to govern fermentation metabolism, sporulation and potentially other important aspects of the organism’s life cycle.

## MATERIALS AND METHODS

### Bacterial strains and media

Bacterial strains utilised in this study are listed in Table S2*. C. acetobutylicum* ATCC 824 and its mutant derivatives were grown at 37°C in an anaerobic cabinet (MG1000 Anaerobic Work Station, Don Whitley Scientific) containing an atmosphere of 80% nitrogen, 10% hydrogen and 10% carbon dioxide. The organism was routinely cultured in supplemented clostridial basal medium (CBMS) (44), unless stated otherwise. CBMS was based on CBM as previously described (45) but contained glucose (50 g/l) and calcium carbonate (5 g/l) as a buffering agent. For agar plates, 10 g/l agar was added and calcium carbonate was omitted. *Escherichia coli* TOP10 was grown in Lysogeny broth at 37°C. Antibiotics were used at the following concentrations: chloramphenicol, 25 μg/ml; erythromycin, 40 μg/ml; tetracycline, 10 μg/ml; thiamphenicol, 15 μg/ml. *C. acetobutylicum* wild type and all mutants generated in this study were stored as spore stocks.

### Plasmids, oligonucleotides, DNA techniques

Plasmids and oligonucleotides used in this study are listed in Tables S3 and S4 (supplemental material) and were synthesised by Eurofins MWG Operon, Germany. PCR amplifications were carried out using high fidelity Phusion polymerase or *Taq* DNA polymerase (both from New England Biolabs). Electroporation of *C. acetobutylicum* was performed as described previously (14). Plasmid isolation and genomic DNA preparations were carried out using the QIAprep Miniprep kit (Qiagen, UK) and DNeasy Blood & Tissue kit (Qiagen), respectively. Restriction enzymes were supplied by New England Biolabs and Promega and were used according to the manufacturers’ instructions. Southern blotting and hybridisation was carried out as previously described (44).

### Construction of mutants using ClosTron technology

Mutants were constructed in *C. acetobutylicum* ATCC 824 using retargeted ClosTron plasmids according to Heap *et al*. (29). The plasmids were designed using the ‘intron targeting and design tool’ available on http://www.ClosTron.com/ClosTron2.php and purchased from DNA2.0. Numbers in the plasmid names (Table S3) indicate the retargeting site used which, in the case of RNPP-type genes, was located within the HTH-encoding region. Genomic DNA from putative mutants was subjected to several PCR screens to establish whether the ClosTron-derived group II intron had inserted into the desired gene target. These included (i) primer pairs that annealed on either side of the target site and (ii) individual flanking primers together with a group II intron specific primer (the latter amplifying the intron–exon junctions). The generated PCR fragments were sequenced to obtain definite proof that insertion had occurred at the desired position. Finally, using chromosomal DNA of all mutant, Southern blot analysis was performed to confirm that only single ClosTron insertions had occurred. At least three independent mutants were generated for each gene.

### Generation of complementation and overexpression vectors

To construct the *qsrB* complementation vector pMTL85141-*qsrB*, a 1659 bp fragment containing the *qsrB* gene and a 351 bp 5’ non-coding region expected to contain the gene’s native promoter were PCR amplified from genomic *C. acetobutylicum* ATCC 824 DNA using the primer pair QsrB_C_F1/QsrB_C_R1 (Table S4). These contained SbfI and NotI restriction sites, respectively, so that the resulting fragment could be cloned into the equally digested clostridial shuttle vector pMTL85141 (32). The resulting vector pMTL85141-*qsrB* was confirmed by restriction analysis and sequencing.

To obtain an overexpression vector in which *qsrB* expression was driven by the strong *C. sporogenes fdx*-promoter the 1336 bp *qsrB* gene was PCR amplified from genomic DNA with the primer pair QsrB_C_F2/QrB_C_R2 (Table S4). These primers contained NdeI and BamHI restriction sites, respectively, which were used to clone the obtained DNA fragment into the clostridial shuttle vector pMTL85143 downstream of the *fdx*-promoter (Dr. Ying Zhang, University of Nottingham, unpublished). The resulting vector pMTL85143-*qsrB* was confirmed by restriction analysis and sequencing.

To obtain the *qspB* expression vector pMTL85143-*qspB*, the 178 bp *qspB* gene was PCR amplified from genomic DNA of *C. acetobutylicum* ATCC 824 using the primer pair QspB_C_F1/QspB_C_R1. These primers contained NdeI and EcoRI restriction sites, respectively, which were used to clone the obtained DNA fragment into the clostridial shuttle vector pMTL85143 downstream of the *fdx*-promoter. The resulting vector pMTL85143-*qspB* was confirmed by restriction analysis and sequencing.

### Spore assays and detection of granulose

*C. acetobutylicum* strains were grown in 5 ml CBMS to enable sporulation. After 7 days, a 200 μl sample of culture was heated to 80°C for 10 min. Serial dilutions were carried out and 20 μl aliquots of the heat-treated cell suspension plated onto CBM agar. Agar plates were incubated for 48 h before CFUs were enumerated. For each assay, a *spo0A* mutant (25) and the wild type were included as negative and positive controls, respectively.

To assess the accumulation of granulose, *C. acetobutylicum* strains were grown on CBM agar containing 5% glucose. Colonies were stained with iodine as previously described (14).

### Determination of colony size

Overnight cultures were serial diluted before plating onto CGM plates (clostridial growth medium containing 1.5% agar; 46) and further incubation for 24 h. To avoid negative impacts on growth, the CGM plates did not contain antibiotics. Measurements were taken from enlarged plate images alongside a scale. For each colony three independent diameter readings were taken and averaged to account for the fact that some colonies were noncircular. A total of twenty colonies per strain were analysed.

### Addition of synthetic QspB fragments to *qsrB* overxpressing strains

Synthetic linear peptides representing C-terminal fragments of the QspB sequence were synthesised and purified by Peptide Protein Research Ltd (Fareham, UK). Thirteen variants were initially obtained (Fig. S5 in supplementary materials) the purity of which was estimated to range from 89-99%, apart from peptides TRSLLGAE, LGAEPTWGWNISKLLF, and TRSLLGAEPTWGWNISKLLF (72%, 79% and 83%, respectively). A selection of peptides showing the highest activities in an initial screen as well as several additional variants (as listed in Fig. 6) were then re-synthesised to >95% purity. Lyophilised peptides were dissolved in DMSO to obtain 20 mM stock solutions. 200 ml of CBMS was inoculated to OD 0.05 with a *C. acetobutylicum* pMTL85143-*qsrB* pre-culture and grown for 4 h. At this point, 10-ml aliquots of the culture were distributed into individual 15-ml Falcon tubes, each containing 5 μl of a particular 20 mM peptide stock solution. Controls only contained 5 μl DMSO. Each peptide or control culture was set up in triplicate.

### Analysis of fermentation products

*C. actetobuytlicum* ATCC 824 wild type and mutants were grown in 50-ml-Falcon tubes containing 30 ml of CBMS. At relevant time points, 1 ml samples were removed, placed on ice and centrifuged at 16,000 x g for 5 min to obtain cells-free culture supernatant. Extraction of fermentation products and their gas chromatographic analysis was carried out as described previously (44).

### Statistical analysis

All numerical data were stored and analysed in IBM SPSS Statistics 19 and 20 (IBM Corporation, Armonk, US) and Microsoft Excel 2007 and 2010. Significance levels were determined with an independent sample two-way t-test in IBM SPSS Statistics (IBM). Data were graphically visualised in GraphPad Prism5 (GraphPad Software, La Jolla, USA) and Excel. Errors bars provided indicate standard deviation.

## ACKNOWLEDGEMENTS

This work was supported by the European Union Marie Curie Initial Training Network ‘CLOSNET’ (contract number 237942) and the Biotechnology and Biological Sciences Research Council (BBSRC; grant numbers BB/G016224/1, BB/J014508/1 and BB/L013940/1). We thank The University of Nottingham for supporting the PhD studentships of OS and ZB.

## REFERENCES

1. Dürre P. 2005. Formation of solvents in clostridia, p671–693. 2005. *In* P. Dürre (ed.), Handbook on clostridia. CRC Press, Boca Raton, Fla.

2. Jones DT, Woods DR. 1986. Acetone-butanol fermentation revisited. Microbiol. Rev. 50:484–524.

3. Jones DT. 2001. Applied acetone-butanol fermentation, p. 125–168. *In* H. Bahl and P. Dürre (ed.), Clostridia. Biotechnology and medical applications. Wiley-VCH Verlag GmbH, Weinheim, Germany.

4. Yoo M, Nguyen N-P-T, Soucaille. 2020. Trends in systems biology for the analysis and Engineering of *Clostridium acetobutylicum* metabolism. Trends Microbiol. 28:118–140.

5. Lütke-Eversloh T, Bahl H. 2011. Metabolic engineering of Clostridium *acetobutylicum*: Recent advances to improve butanol production. Curr. Opin. Biotech. 22:1–14.

6. Xue C, Cheng C. 2019. Chapter 2 –Butanol production by *Clostridium*, Adv. Bioenerg. 4: 35–77.

7. Ravagnani, A, Jennert KC, Steiner, Grünberg ER, Jefferies JR, Wilkinson SR, Young DI, Tidswell EC, Brown DP, Youngman P, Morris JG, and Young M. 2000. Spo0A directly controls the switch from acid to solvent production in solvent-forming clostridia. Mol. Microbiol. 37:1172–1185.

8. Harris LM, Welker NE, Papoutsakis ET. 2002. Northern, morphological, and fermentation analysis of *spo0A* inactivation and overexpression in *Clostridium acetobutylicum* ATCC 824. J. Bacteriol. 184:3586–3597.

9. Thormann K, Feustel L, Lorenz K, Nakotte S, Durre P. 2002. Control of butanol formation in *Clostridium acetobutylicum* by transcriptional activation. J Bacteriol 184:1966–1973.

10. Nold, N. 2008. Untersuchungen zur Regulation des sol-Operons in Clostridium acetobutylicum. PhD thesis, University of Ulm.

11. Zimmermann, T. 2013. Untersuchungen zur Butanolbildung von Hyperthermus butylicus und Clostridium acetobutylicum. PhD thesis, University of Ulm.

12. Ren C, Gu Y, Wu Y, Zhang W, Yang C, Yang S, Jiang W. 2012. Pleiotropic functions of catabolite control protein CcpA in butanol-producing *Clostridium acetobutylicum*. BMC Genomics 13:349.

13. Jones AJ, Fast A. G, Clupper M, Papoutsakis ET. 2018. Small and low but potent: the complex regulatory role of the small RNA SolB in solventogenesis in *Clostridium acetobutylicum*. Appl. Environ. Microbiol. 84:e00597–18,

14. Steiner E, Dago AE, Young DI, Heap JT, Minton NP, Hoch JA, Young M. 2011. Multiple orphan histidine kinases interact directly with Spo0A to control the initiation of endospore formation in *Clostridium acetobutylicum*. Mol. Microbiol. 80:641–654.

15. Zhao Y, Tomas CA, Rudolph FB, Papoutsakis ET, Bennett GN. 2005. Intracellular butyryl phosphate and acetyl phosphate concentrations in *Clostridium acetobutylicum* and their implications for solvent formation. Appl. Environ. Microbiol. 71:530–537.

16. Steiner E, Scott J, Minton NP, Winzer K. 2012. An agr quorum sensing system that regulates granulose formation and sporulation in *Clostridium acetobutylicum*. Appl. Environ. Microbiol. 78:1113–1122.

17. Sturme MHJ, Kleerebezem M, Nakayama J, Akkermans ADL, Vaughan EE, de Vos WM. 2002. Cell to cell communication by autoinducing peptides in Gram-positive bacteria. Antonie Van Leeuwenhoek 81:233–243.

18. Lyon GJ, Novick RP. 2004. Peptide signaling in *Staphylococcus aureus* and other Gram-positive bacteria. Peptides 25:1389–1403.

19. Kosaka T, Nakayama S, Nakaya K, Yoshino S, Furukawa K. 2007. Characterization of the sol operon in butanol-hyperproducing *Clostridium saccharoperbutylacetonicum* strain N1-4 and its degeneration mechanism. Biosci. Biotechnol. Biochem. 71:58–68.

20. Wuster A, Babu MM. 2008. Conservation and evolutionary dynamics of the agr cell-to-cell communication system across firmicutes. J. Bacteriol. 190:743–746.

21. Herman N, Kim S-J, Li JS, Cai W, Koshino H, Zhang W. 2017. The industrial anaerobe *Clostridium acetobutylicum* uses polyketides to regulate butanol production and differentiation. Nat. Commun. 15:1514.

22. Pottathil M, Lazazzera BA. 2003. The extracellular Phr peptide-Rap phosphatase signaling circuit of *Bacillus subtilis*. Front. Biosci. 8:32–45.

23. Declerck N, Bouillaut L, Chaix D, Rugani N, Slamti L, Hoh F, Lereclus D, Arold ST. 2007. Structure of PlcR: Insights into virulence regulation and evolution of quorum sensing in Gram-positive bacteria. Proc. Natl. Acad. Sci. USA. 104:18490–18495.

24. Rocha-Estrada J, Aceves-Diez A, Guarneros G, De La Torre M. 2010. The RNPP family of quorum-sensing proteins in Gram-positive bacteria. Appl. Microbiol. Biotechnol. 87:913–923.

25. Ehsaan M, Kuit W, Zhang Y, Cartman ST, Heap JT, Winzer K, Minton NP. 2016. Mutant generation by allelic exchange and genome resequencing of the biobutanol organism *Clostridium acetobutylicum* ATCC 824. Biotechnol. Biofuels 9:4.

26. Hu S, Zheng H, Gu Y, Zhao J, Zhang W, Yang Y, Wang S, Zhao G, Yang S, Jiang W. 2011. Comparative genomic and transcriptomic analysis revealed genetic characteristics related to solvent formation and xylose utilization in *Clostridium acetobutylicum* EA 2018. BMC Genomics 12:93

27. Bao G, Wang R, Zhu Y, Dong H, Mao S, Zhang Y, Chen Z, Li Y, Ma Y. 2011. Complete genome sequence of *Clostridium acetobutylicum* DSM 1731, a solvent-producing strain with multireplicon genome architecture. J. Bacteriol. 193:5007–5008.

28. Karpenahalli M, Lupas A, Soding, J. 2007. TPRpred: a tool for prediction of TPR-, PPR- and SEL1-like repeats from protein sequences. BMC Bioinformatics 8:2.

29. Heap JT, Kuehne SA, Ehsaan M, Cartman ST, Cooksley CM, Scott JC, Minton NP. 2010. The ClosTron: mutagenesis in Clostridium refined and streamlined. J. Microbiol. Methods 80:49–55.

30. Hartmanis MGN, Ahlman H, Gatenbeck S. 1986. Stability of solvent formation in Clostridium acetobutylicum during repeated subculturing. Appl. Microbiol. Biotechnol. 23:369–371.

31. Kashket ER, Cao Z-Y. 1995. Clostridial strain degeneration. FEMS Microbiol Rev. 17:307–316.

32. Heap JT, Pennington OJ, Cartman ST, Minton NP. 2009. A modular system for Clostridium shuttle plasmids. J. Microbiol. Methods 78:79–85.

33. Gohar M, Faegri K, Perchat S, Ravnum S, Okstad OA, Gominet M, Kolsto AB, Lereclus D. 2008. The PlcR virulence regulon of Bacillus cereus. PLoS ONE 3:e2793.

34. Maddox IS, Steiner E, Hirsch S, Wessner S, Gutierrez NA, Gapes JR, Schuster KC. 2000. The cause of “acid-crash” and “acidogenic fermentations” during the batch acetone-butanol-ethanol (ABE-) fermentation process. J. Mol. Microbiol. Biotechnol. 2:95–100.

35. Diggle SP, West SA, Gardner A, Griffin AS. 2008. Communication in bacteria. In Sociobiology of communication: An interdisciplinary perspective (ed. David Hughes & Patrizia D’Ettorre): pp. 11–31. Oxford University Press.

36. Kalamara M, Spacapan M, Mandic-Mulec I, Stanley-Wall NR. 2018. Social behaviours by *Bacillus subtilis*: quorum sensing, kin discrimination and beyond. Mol. Microbiol. 110:863–878

37. Poehlein A, Solano JDM, Flitsch SK, Krabben P, Winzer K, Reid SJ, Joes DT, Green E, Minton NP, Daniel R, Dürre P. 2017. Microbial solvent formation revisited by comparative genome analysis. Biotechnology for Biofuels. 10:58

38. Jones DT, Webster JR, Woods DR. 1980. The formation of simple fruiting body like structures associated with sporulation under aerobic conditions. J. Gen. Microbiol. 116:195–200.

39. Cornforth DM, Popat R, McNally L, Gurney J, Scott-Phillips TC, Ivens A, Diggle SP, Brown SP. 2014 Combinatorial quorum sensing allows bacteria to resolve their social and physical environment. Proc. Natl Acad. Sci. USA 111:4280–4284.

40. Kotte A-K. 2013. RNPP-type Quorum Sensing in Clostridium acetobutylicum. PhD thesis, University of Nottingham.

41. Edwards AN, Tamayo R, McBride SMA. 2016. Novel regulator controls *Clostridium difficile* sporulation, motility and toxin production. Mol. Microbiol. 100:954–971.

42. Bouillaut L, Perchat S, Arold S, Zorrilla S, Slamti L, Henry C, Gohar M, Declerck N, Lereclus D. 2008. Molecular basis for group-specific activation of the virulence regulator PlcR by PapR heptapeptides. Nucleic Acids Res. 36:3791–3801.

43. Slamti L, Lereclus D. 2005. Specificity and polymorphism of the PlcR-PapR quorum-sensing system in the Bacillus cereus group. J. Bacteriol. 187:1182–1187.

44. Cooksley CM, Zhang Y, Wang H, Redl S, Winzer K, Minton NP. 2012. Targeted mutagenesis of the Clostridium acetobutylicum acetone–butanol–ethanol fermentation pathway. Metab. Eng. 14:630–641.

45. O’Brien RW, Morris JG. 1971. Oxygen and the growth and metabolism of *Clostridium acetobutylicum*. J. Gen. Microbiol. 68:307–318.

46. Hartmanis MGN, Gatenbeck S. 1984. Intermediary metabolism in *Clostridium acetobutylicum*: levels of enzymes involved in the formation of acetate and butyrate. Appl. Environ. Microbiol. 47:1277–1283.

