## Supplemental Figure S1 for "RNPP-type quorum sensing regulates solvent formation and sporulation in *Clostridium acetobutylicum*"

### Slide 1
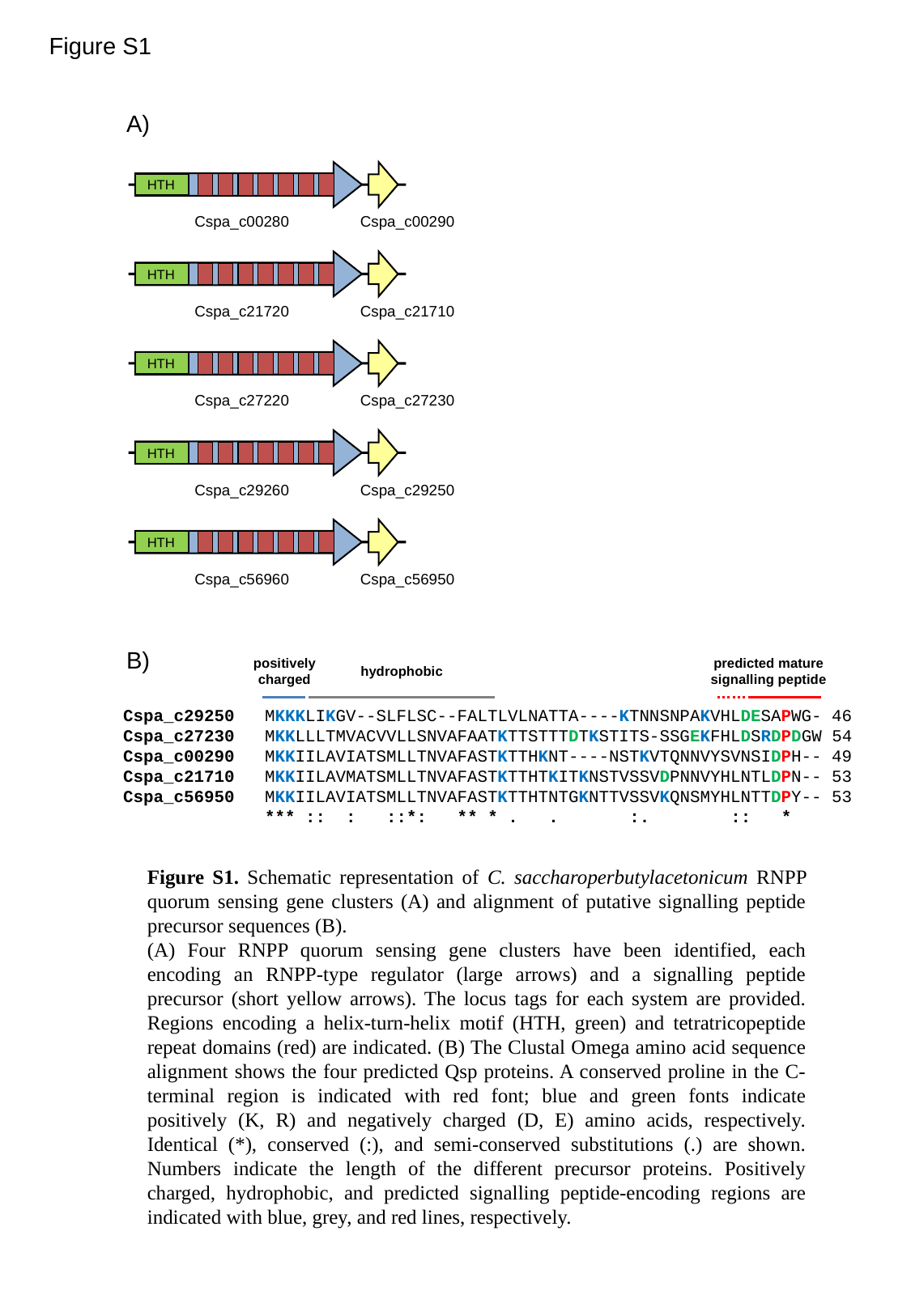

Figure S1
A)
HTH
Cspa_c00280
Cspa_c00290
HTH
Cspa_c21720
Cspa_c21710
HTH
Cspa_c27220
Cspa_c27230
HTH
Cspa_c29260
Cspa_c29250
HTH
Cspa_c56960
Cspa_c56950
B)
positively
charged
predicted mature
signalling peptide
hydrophobic
Cspa_c29250 MKKKLIKGV--SLFLSC--FALTLVLNATTA----KTNNSNPAKVHLDESAPWG- 46
Cspa_c27230 MKKLLLTMVACVVLLSNVAFAATKTTSTTTDTKSTITS-SSGEKFHLDSRDPDGW 54
Cspa_c00290 MKKIILAVIATSMLLTNVAFASTKTTHKNT----NSTKVTQNNVYSVNSIDPH-- 49
Cspa_c21710 MKKIILAVMATSMLLTNVAFASTKTTHTKITKNSTVSSVDPNNVYHLNTLDPN-- 53
Cspa_c56950 MKKIILAVIATSMLLTNVAFASTKTTHTNTGKNTTVSSVKQNSMYHLNTTDPY-- 53
 *** :: : ::*: ** * . . :. :: *
Figure S1. Schematic representation of C. saccharoperbutylacetonicum RNPP quorum sensing gene clusters (A) and alignment of putative signalling peptide precursor sequences (B).
(A) Four RNPP quorum sensing gene clusters have been identified, each encoding an RNPP-type regulator (large arrows) and a signalling peptide precursor (short yellow arrows). The locus tags for each system are provided. Regions encoding a helix-turn-helix motif (HTH, green) and tetratricopeptide repeat domains (red) are indicated. (B) The Clustal Omega amino acid sequence alignment shows the four predicted Qsp proteins. A conserved proline in the C-terminal region is indicated with red font; blue and green fonts indicate positively (K, R) and negatively charged (D, E) amino acids, respectively. Identical (*), conserved (:), and semi-conserved substitutions (.) are shown. Numbers indicate the length of the different precursor proteins. Positively charged, hydrophobic, and predicted signalling peptide-encoding regions are indicated with blue, grey, and red lines, respectively.
