## Supplemental Figure S2 for "RNPP-type quorum sensing regulates solvent formation and sporulation in *Clostridium acetobutylicum*"

**B)**

**A)**

|  |  |  |  |  |  |
| --- | --- | --- | --- | --- | --- |
| 0 | 0.0575 | 0.0575 | 0.065 | 0.06 | 0.06 |
| 3 | 0.285 | 0.2175 | 0.255 | 0.1675 | 0.3325 |
| 6 | 1.72 | 1.4375 | 1.1175 | 0.9375 | 2.15875 |
| 9 | 2.9725 | 2.125 | 2.3625 | 2.4425 | 2.59625 |
| 12 | 2.485 | 1.305 | 2.345 | 3.125 | 2.4925 |
| 24 | 1.41 | 0.9875 | 1.5875 | 1.7625 | 1.23 |
| 48 | 3.2375 | 3.4325 | 2.1925 | 3.01 | 2.885 |

|  |  |  |  |  |  |
| --- | --- | --- | --- | --- | --- |
| 0 | 0.06 | 0.0675 | 0.0633333 | 0.06 | 0.0566667 |
| 3 | 0.3325 | 0.38 | 0.406667 | 0.28 | 0.203333 |
| 6 | 2.15875 | 2.1975 | 1.54 | 1.14667 | 1.25333 |
| 9 | 2.59625 | 2.895 | 2.35333 | 2.10667 | 2.42333 |
| 12 | 2.4925 | 2.02 | 2.40667 | 2.06667 | 2.23333 |
| 24 | 1.23 | 0.935 | 1.25 | 1.38667 | 1.42667 |
| 48 | 2.885 | 2.91 | 1.91667 | 1.02667 | 2.10333 |

**Figure S2.** Growth of *C. acetobutylicum* ATCC 824 and derived *qsr* mutants in CBMS medium.

The data represent the mean of three independently cultures. A) Wild type (closed circles); *qsrA* (open squares), *qsrB* (open circles), *qsrC* (closed triangles) and *qsrD* (open diamonds) mutants. B) Wild type (closed circles); *qsrE* (open squares), *qsrF* (open circles), *qsrG* (closed triangles) and *qsrH* (open diamonds) mutants.
