## Supplemental Figure S3 for "RNPP-type quorum sensing regulates solvent formation and sporulation in *Clostridium acetobutylicum*"

### Slide 1
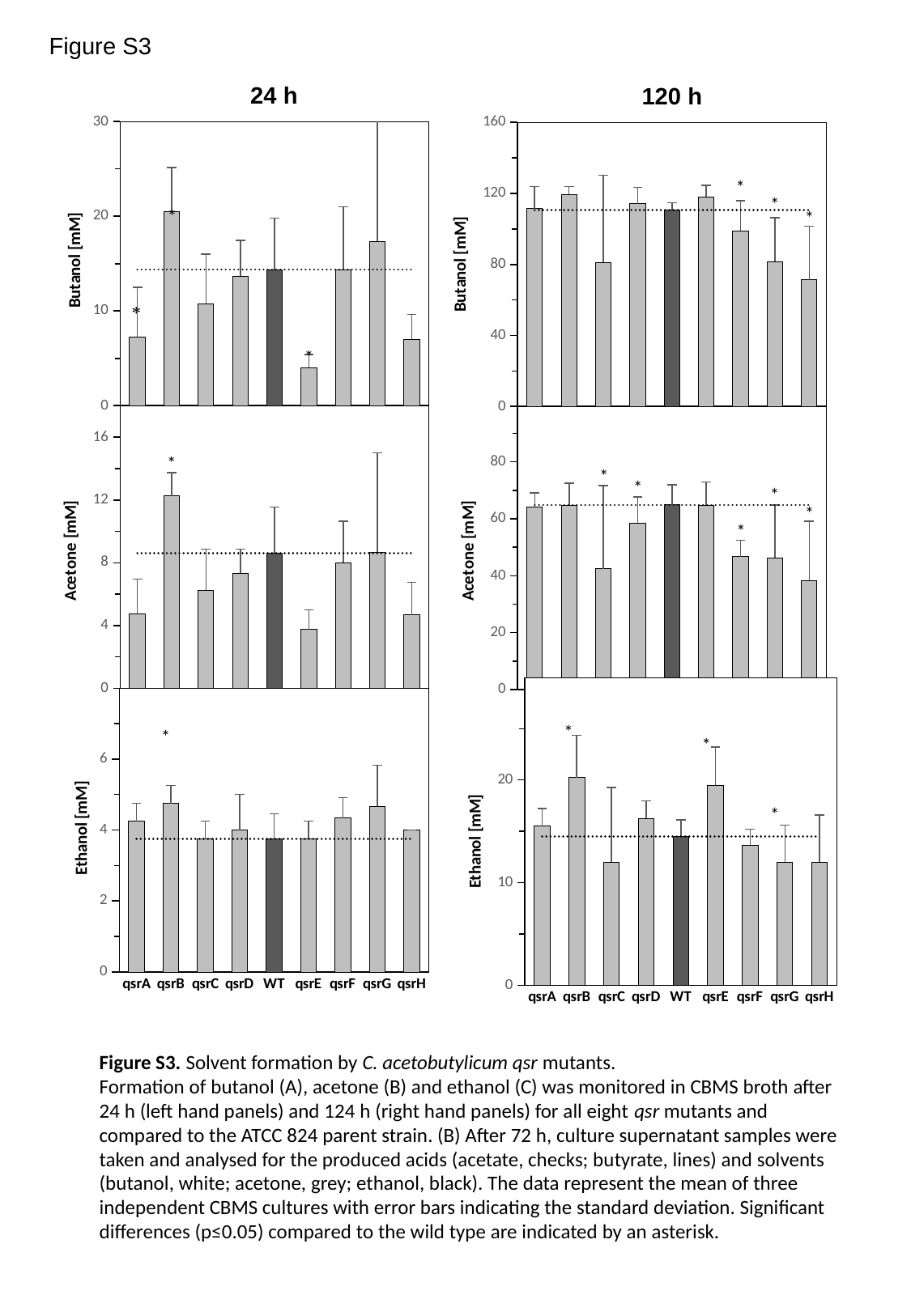

Figure S3
24 h
#### Chart
| Category | | Reference lines |
|---|---|---|
| qsrA | 7.25 | 14.375 |
| qsrB | 20.5 | 14.375 |
| qsrC | 10.75 | 14.375 |
| qsrD | 13.666666666666666 | 14.375 |
| WT | 14.375 | 14.375 |
| qsrE | 4.0 | 14.375 |
| qsrF | 14.333333333333334 | 14.375 |
| qsrG | 17.333333333333332 | 14.375 |
| qsrH | 7.0 | 14.375 |
#### Chart
| Category | | Reference lines |
|---|---|---|
| qsrA | 4.75 | 8.625 |
| qsrB | 12.25 | 8.625 |
| qsrC | 6.25 | 8.625 |
| qsrD | 7.333333333333333 | 8.625 |
| WT | 8.625 | 8.625 |
| qsrE | 3.75 | 8.625 |
| qsrF | 8.0 | 8.625 |
| qsrG | 8.666666666666666 | 8.625 |
| qsrH | 4.666666666666667 | 8.625 |
#### Chart
| Category | | Reference lines |
|---|---|---|
| qsrA | 4.25 | 3.75 |
| qsrB | 4.75 | 3.75 |
| qsrC | 3.75 | 3.75 |
| qsrD | 4.0 | 3.75 |
| WT | 3.75 | 3.75 |
| qsrE | 3.75 | 3.75 |
| qsrF | 4.333333333333333 | 3.75 |
| qsrG | 4.666666666666667 | 3.75 |
| qsrH | 4.0 | 3.75 |120 h
#### Chart
| Category | | Reference lines |
|---|---|---|
| qsrA | 111.5 | 110.75 |
| qsrB | 119.5 | 110.75 |
| qsrC | 81.25 | 110.75 |
| qsrD | 114.25 | 110.75 |
| WT | 110.75 | 110.75 |
| qsrE | 118.0 | 110.75 |
| qsrF | 99.0 | 110.75 |
| qsrG | 81.66666666666667 | 110.75 |
| qsrH | 71.33333333333333 | 110.75 |
#### Chart
| Category | | Reference lines |
|---|---|---|
| qsrA | 64.25 | 64.875 |
| qsrB | 64.75 | 64.875 |
| qsrC | 42.5 | 64.875 |
| qsrD | 58.5 | 64.875 |
| WT | 64.875 | 64.875 |
| qsrE | 64.75 | 64.875 |
| qsrF | 46.666666666666664 | 64.875 |
| qsrG | 46.333333333333336 | 64.875 |
| qsrH | 38.333333333333336 | 64.875 |
#### Chart
| Category | | Reference lines |
|---|---|---|
| qsrA | 15.5 | 14.5 |
| qsrB | 20.25 | 14.5 |
| qsrC | 12.0 | 14.5 |
| qsrD | 16.25 | 14.5 |
| WT | 14.5 | 14.5 |
| qsrE | 19.5 | 14.5 |
| qsrF | 13.666666666666666 | 14.5 |
| qsrG | 12.0 | 14.5 |
| qsrH | 12.0 | 14.5 |
Figure S3. Solvent formation by C. acetobutylicum qsr mutants.
Formation of butanol (A), acetone (B) and ethanol (C) was monitored in CBMS broth after 24 h (left hand panels) and 124 h (right hand panels) for all eight qsr mutants and compared to the ATCC 824 parent strain. (B) After 72 h, culture supernatant samples were taken and analysed for the produced acids (acetate, checks; butyrate, lines) and solvents (butanol, white; acetone, grey; ethanol, black). The data represent the mean of three independent CBMS cultures with error bars indicating the standard deviation. Significant differences (p≤0.05) compared to the wild type are indicated by an asterisk.
