## Supplemental Figure S4 for "RNPP-type quorum sensing regulates solvent formation and sporulation in *Clostridium acetobutylicum*"

Solvents [mM]

| Time [h] | WT | qsrB | WT | qsrB | WT | qsrB |
| --- | --- | --- | --- | --- | --- | --- |
| 0 | 1.125 | 1 | 0.75 | 1 | 1.375 | 1.75 |
| 9 | 3.375 | 2.75 | 2.5 | 3.75 | 2.125 | 3 |
| 12 | 2.625 | 2.75 | 3.375 | 6.75 | 3.25 | 4.5 |
| 24 | 3.75 | 4.75 | 14.375 | 20.5 | 8.625 | 12.25 |
| 48 | 6.5 | 7.75 | 34.25 | 53.25 | 18.75 | 31.75 |
| 72 | 9.25 | 10.5 | 60.5 | 76.75 | 34.125 | 43 |
| 96 | 11.25 | 15 | 87.25 | 99.5 | 50.5 | 54.5 |
| 120 | 14.5 | 20.25 | 110.75 | 119.5 | 64.875 | 64.75 |

Butyrate/Acetone [mM]

| Time [h] | WT | qsrB | WT | qsrB |
| --- | --- | --- | --- | --- |
| 0 | 0.5 | 0.5 | 0.25 | 0.25 |
| 9 | 5.5 | 4.25 | 16 | 11.5 |
| 12 | 6.75 | 6.25 | 19.25 | 14.25 |
| 24 | 8.5 | 7.25 | 22 | 14 |
| 48 | 8.75 | 5.75 | 52.125 | 22.75 |
| 72 | 7.25 | 5 | 31.75 | 14.25 |
| 96 | 6.625 | 6 | 8.75 | 4 |
| 120 | 7.125 | 7 | 10.25 | 3.25 |

**Figure S4.** Solvent and acid production by *C. acetobutylicum* *qsrB* mutants

(A) Concentration of butanol (circles), acetone (squares) and ethanol (triangles) in the culture supernatant at the indicated time points. Open and closed symbols represent *qsrB* mutant and ATTC 824 parent strain data, respectively. (B) Concentration of butyrate (circles) and acetate (squares) in the culture supernatant at the indicated time point. Open and closed symbols represent *qsrB* mutant and ATTC 824 parent strain data, respectively. Data represent the mean of three independent cultures with error bars indicating the standard deviation. Significant differences (p≤ 0.05) compared to the wild type are indicated by an asterisk next to the relevant data point.
