## Supplemental Figure S5 for "RNPP-type quorum sensing regulates solvent formation and sporulation in *Clostridium acetobutylicum*"

### Slide 1
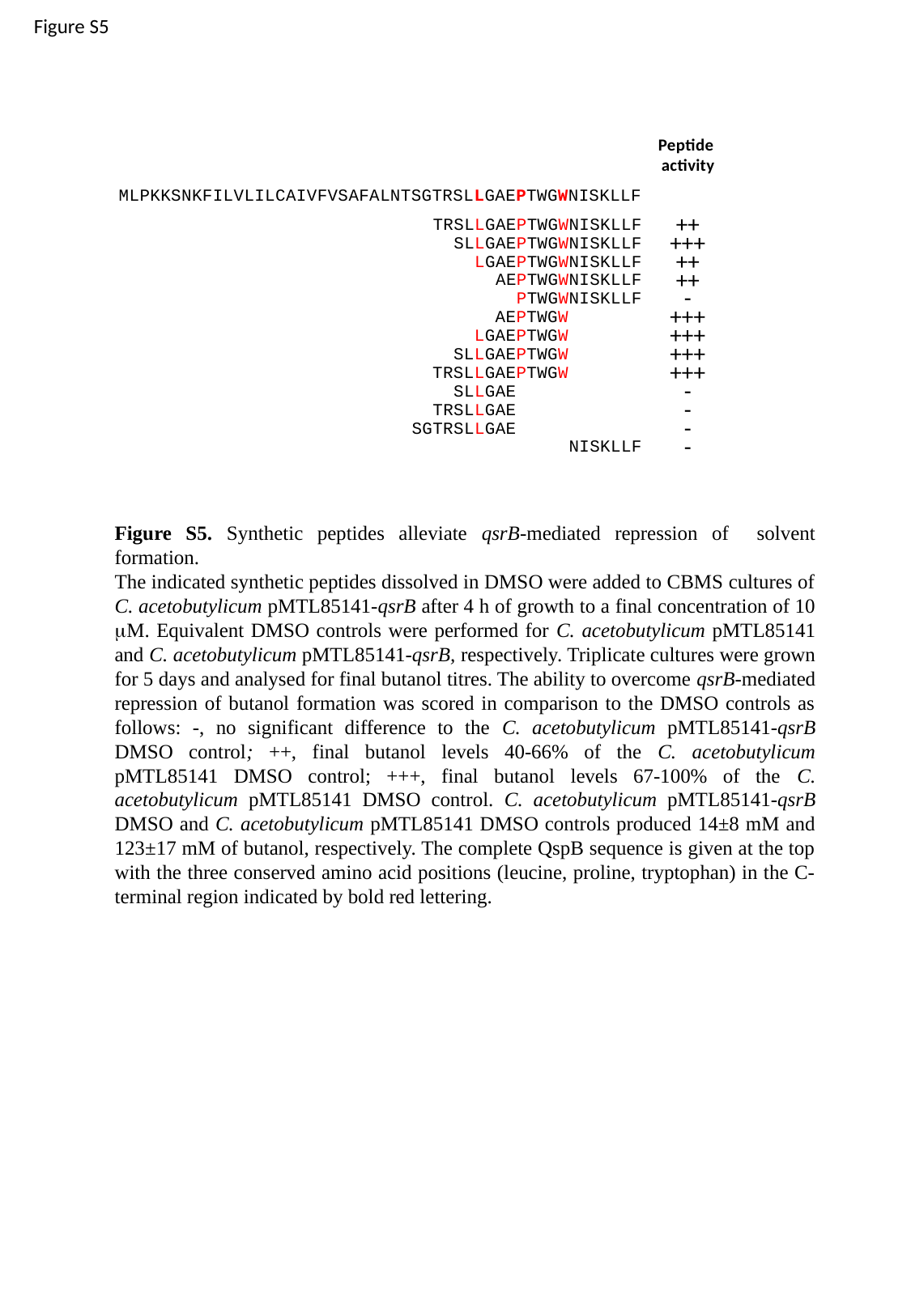

Figure S5
Peptide
activity
MLPKKSNKFILVLILCAIVFVSAFALNTSGTRSLLGAEPTWGWNISKLLF
++
TRSLLGAEPTWGWNISKLLF
+++
SLLGAEPTWGWNISKLLF
++
LGAEPTWGWNISKLLF
++
AEPTWGWNISKLLF
-
PTWGWNISKLLF
+++
AEPTWGW
+++
LGAEPTWGW
+++
SLLGAEPTWGW
+++
TRSLLGAEPTWGW
-
SLLGAE
-
TRSLLGAE
-
SGTRSLLGAE
-
NISKLLF
Figure S5. Synthetic peptides alleviate qsrB-mediated repression of solvent formation.
The indicated synthetic peptides dissolved in DMSO were added to CBMS cultures of C. acetobutylicum pMTL85141-qsrB after 4 h of growth to a final concentration of 10 μM. Equivalent DMSO controls were performed for C. acetobutylicum pMTL85141 and C. acetobutylicum pMTL85141-qsrB, respectively. Triplicate cultures were grown for 5 days and analysed for final butanol titres. The ability to overcome qsrB-mediated repression of butanol formation was scored in comparison to the DMSO controls as follows: -, no significant difference to the C. acetobutylicum pMTL85141-qsrB DMSO control; ++, final butanol levels 40-66% of the C. acetobutylicum pMTL85141 DMSO control; +++, final butanol levels 67-100% of the C. acetobutylicum pMTL85141 DMSO control. C. acetobutylicum pMTL85141-qsrB DMSO and C. acetobutylicum pMTL85141 DMSO controls produced 14±8 mM and 123±17 mM of butanol, respectively. The complete QspB sequence is given at the top with the three conserved amino acid positions (leucine, proline, tryptophan) in the C-terminal region indicated by bold red lettering.
