## Supplemental Table S1 for "RNPP-type quorum sensing regulates solvent formation and sporulation in *Clostridium acetobutylicum*"

TABLE S1. Formation of heat resistant endospores by *qsr* mutants

| Strain | Heat-resistant CFU/ml | p^1^ |
| --- | --- | --- |
| *C. acetobutylicum qsrA*::CT*ermB* | 9.50 × 10^7^ | 0.561 |
| *C. acetobutylicum qsrB*::CT*ermB* | 1.18 × 10^8^ | 0.929 |
| *C. acetobutylicum qsrC*::CT*ermB* | 7.57 × 10^7^ | 0.323 |
| *C. acetobutylicum qsrD*::CT*ermB* | 8.91 × 10^7^ | 0.473 |
| *C. acetobutylicum qsrE*::CT*ermB* | 1.26 × 10^8^ | 0.743 |
| *C. acetobutylicum qsrF*::CT*ermB* | 8.22 × 10^7^ | 0.339 |
| *C. acetobutylicum qsrG*::CT*ermB* | 3.85 × 10^7^ | *0.036 |
| *C. acetobutylicum qsrH*::CT*ermB* | 1.21 × 10^8^ | 0.857 |
| *C.* acetobutylicum ATCC 824 | 1.15 × 10^8^ | - |
| ^1^Significant differences to the wild type are indicated by an asterisk. | | |
