## Supplemental Table S2 for "RNPP-type quorum sensing regulates solvent formation and sporulation in *Clostridium acetobutylicum*"

Table S2. Bacterial strains used in this study

| **Strain** | **Relevant properties** | **Source/reference** |
| --- | --- | --- |
| *E. coli* Top10 | F- *mcrA* Δ(*mrr-hsdRMS-mcrBC*)  Φ80*lacZ*ΔM15 Δ*lacX*74 *recA*1  *araD*139 Δ(*ara leu*)7697 *galU* *galK rpsL* (StrR) endA1 nupG | Invitrogen |
| *E. coli* Top 10 pAN2 | *E. coli* Top 10 with methylation plasmid pAN2 containing the ϕ3TI methyltransferase | Heap et al. (2007) |
| *C. acetobutylicum* ATCC 824 | *C. acetobutylicum* ATCC 824 wild type | Prof. Hubert Bahl, University of Rostock (COSMIC-strain) |
| *C. acetobutylicum* *qsrA*::CT*ermB* | *C. acetobutylicum* ATCC 824 *qsrA* ClosTron mutant | This work |
| *C. acetobutylicum* *qsrB*::CT*ermB* | *C. acetobutylicum* ATCC 824 *qsrB* ClosTron mutant | This work |
| *C. acetobutylicum* *qsrC*::CT*ermB* | *C. acetobutylicum* ATCC 824 *qsrC* ClosTron mutant | This work |
| *C. acetobutylicum* *qsrD*::CT*ermB* | *C. acetobutylicum* ATCC 824 *qsrD* ClosTron mutant | This work |
| *C. acetobutylicum* *qsrE*::CT*ermB* | *C. acetobutylicum* ATCC 824 *qsrE* ClosTron mutant | This work |
| *C. acetobutylicum* *qsrF*::CT*ermB* | *C. acetobutylicum* ATCC 824 *qsrF* ClosTron mutant | This work |
| *C. acetobutylicum* *qsrG*::CT*ermB* | *C. acetobutylicum* ATCC 824 *qsrG* ClosTron mutant | This work |
| *C. acetobutylicum* *qsrH*::CT*ermB* | *C. acetobutylicum* ATCC 824 *qsrH* ClosTron mutant | This work |
| *C. acetobutylicum* *qspB*::CT*ermB* | *C. acetobutylicum* ATCC 824 *qspB* ClosTron mutant | This work |
| *C. acetobutylicum* pMTL85141 | ATTC 824 wild type with empty pMTL85141 vector | This work |
| *C. acetobutylicum* pMTL85143 | ATTC 824 wild type with empty pMTL85143 vector | This work |
| *C. acetobutylicum* *qsrB*::CT*ermB* pMTL85141 | *qsrB* mutant with empty ATTC 824 wild type with empty pMTL85141 vector | This work |
| *C. acetobutylicum* *qsrB*::CT*ermB* pMTL85141-*qsrB* | Complemented *qsrB* mutant carrying pMTL85141-*qsrB* | This work |
| *C. acetobutylicum* *qsrB*::CT*ermB* pMTL85143 | *qsrB* mutant with empty pMTL85143 vector | This work |
| *C. acetobutylicum* *qsrB*::CT*ermB* pMTL85143-*qsrB* | Complemented *qsrB* mutant carrying pMTL85143-*qsrB* | This work |
| *C. acetobutylicum* pMTL85141-*qsrB* | *qsrB* overexpressing ATTC 824 wild type carrying pMTL85141-*qsrB* | This work |
| *C. acetobutylicum* pMTL85143-*qsrB* | *qsrB* overexpressing ATTC 824 wild type carrying pMTL85143-*qsrB* | This work |
| *C. acetobutylicum* *qspB*::CT*ermB* pMTL85143 | *qspB* mutant with empty plasmid | This work |
| *C. acetobutylicum* *qspB*::CT*ermB* pMTL85143-qspB | Complemented *qspB* mutant | This work |
| *C. acetobutylicum* pMTL85143-*qspB* | *qspB* overexpressing ATTC 824 wild type carrying pMTL85143-*qssB* | This work |
