## Supplemental Table S3 for "RNPP-type quorum sensing regulates solvent formation and sporulation in *Clostridium acetobutylicum*"

Table S3. Plasmids used in this study

| **Plasmid** | **Relevant properties** | **Source** |
| --- | --- | --- |
| pAN2 | Plasmid containing ϕ3TI methyltransferase | Heap et al. (2007) |
| pCR2.1-TOPO | A plasmid that is supplied linearized with A-overhangs for convenient cloning of PCR fragments | Invitrogen |
| pMTL007C-E2::*qsrA*‑102\|103A | ClosTron plasmid retargeted to *qsrA*^1^ | This study |
| pMTL007C-E2::*qsrB*‑102\|103S | ClosTron plasmid retargeted to *qsrB*^1^ | This study |
| pMTL007C-E2::*qsrC*‑102\|103S | ClosTron plasmid retargeted to *qsrC*^1^ | This study |
| pMTL007C-E2::*qsrD*‑49\|50A | ClosTron plasmid retargeted to *qsrD*^1^ | This study |
| pMTL007C-E2::*qsrES*‑58\|59A | ClosTron plasmid retargeted to *qsrE*^1^ | This study |
| pMTL007C-E2::*qsrF*‑107\|108A | ClosTron plasmid retargeted to *qsrF*^1^ | This study |
| pMTL007C-E2::*qsrG*‑93\|94A | ClosTron plasmid retargeted to *qsrG*^1^ | This study |
| pMTL007C-E2::*qsrH*‑58\|59A | ClosTron plasmid retargeted to *qsrH*^1^ | This study |
| pMTL007C-E2::*qspB*‑53/54A | ClosTron plasmid retargeted to *qspB*^1^ | This study |
| pMTL85141 | Clostridium modular plasmid containing *catP* | Heap et al. (2009) |
| pMTL85143 | pMTL85141 with *C. sporogenes* ferredoxin promoter upstream of multiple cloning site | Dr Ying Zhang,  Univ. of Nottingham |
| pMTL85141-*qsrB* | pMTL85141 containing *qsrB* coding region and 351 bp non-coding region upstream | This study |
| pMTL85143-*qsrB* | pMTL85143 containing *qsrB* coding region | This study |
| pMTL85143-*qspB* | pMTL85143 containing *qspB* coding region | This study |

^1^Numbers following the gene name indicate the predicted insertion site of the encoded ClosTron derivative, with S and A denoting sense and anti-sense orientation, respectively.
