## Supplemental Table S4 for "RNPP-type quorum sensing regulates solvent formation and sporulation in *Clostridium acetobutylicum*"

Table S4. Oligonucleotides used in this study

| **Oligonucleotide** | **Sequence (5’ to 3’)** |
| --- | --- |
| **ClosTron mutant screening** |  |
| QsrA_F | AAGAGGAATTAGCGGGAGCTGAG |
| QsrA_R | CGACTTCTGTCAATTTGGTTGAGAAGC |
| QsrB_F | CGATATTGTTGGAGAAGAAGTTACTC |
| QsrB_R | AGATAATCCGCAGTTACATCC |
| QsrC_F | TCAAATACTGCCTATTGGCGTAAAGC |
| QsrC_R | AGCATTATTTCTGCTGCATGTCTAG |
| QsrD_F | GGAGAGTTTTGTCATATGTGTGTC |
| QsrD_R | AGCTTGTGATTCCTCATCCTC |
| QsrE_F | GATAAGGGAGAAAGTGCTATGGCAAG |
| QsrE_R | TCCTCTTGAAAAGGCATCTCTCTT |
| QsrF_F | AGATGATATTGTAGGTACAGAACTCAC |
| QsrF_R | GTCCTGTATGTATGAGGCGATC |
| QsrG_F | ACGGCCTAAGTCAAGAAGATCTGG |
| QsrG_R | ATTGCTTGCGATTTCTCATCTTCCATC |
| QsrH_F | GCACTTATGAGATAATGTCTATTGGAGACAAGC |
| QsrH_R | TGCTGCACTTCTAGTAAGGTTTGCT |
| EBS universal | CGAAATTAGAAACTTGCGTTCAGTAAAC |
| **Cloning** |  |
| QsrB_C_F1 | TATATACCTGCAGGCTACATTACTCAAAGCATATAAATACG |
| QsrB_ C_R1 | TATATAGCGGCCGCTTACTTAAACTTTATTAAAAAATTTAATATTTTATCTATGTC |
| QsrB_ C_F2 | CTTGGTCATATGGGAAACTGTC |
| QsrB_ C_R2 | AACATCGGATCCTATTTACTTACTTAAAC |
| QspB_ C_F1 | TGTCTACATATGTTACCAAAAAAGAGTAATAAATTTATATTAG |
| QspB_ C_R1 | TTTTTAGAATTCGGTTTTTGTTTAATGTTATAAAAC |
| **Southern Blot probe generation** | |
| EBS2 | TGAACGCAAGTTTCTAATTTCGGTTCTCATCCGATAGAGGAAAGTGTCT |
| Intron SalI-R1 | ATTACTGTGACTGGTTTGCACCACCCTCTTCG |
